# Inducer of CBF Expression 1 (ICE1) Promotes Cold-enhanced Immunity by Directly Activating Salicylic Acid Signaling

**DOI:** 10.1101/2023.09.12.557434

**Authors:** Shaoqin Li, Li He, Yongping Yang, Yixin Zhang, Xiao Han, Yanru Hu, Yanjuan Jiang

**Affiliations:** CAS Key Laboratory of Tropical Plant Resources and Sustainable Use, Xishuangbanna Tropical Botanical Garden, Chinese Academy of Sciences, Kunming, Yunnan 650223, China; University of Chinese Academy of Sciences, Beijing 100049, China; State Key Laboratory for Conservation and Utilization of Bio-Resources in Yunnan, School of Life Sciences, Yunnan University, Kunming 650091, China

**Author notes:** The author(s) responsible for distribution of materials integral to the findings presented in this article in accordance with the policy described in the Instructions for Authors is (are): Yanjuan Jiang.

## Abstract

Cold stress affects plant immune responses, and this process may involve the salicylic acid (SA) signaling pathway. However, the underlying mechanism by which low temperature signals coordinate with SA signaling to regulate plant immunity remains not fully understood. Here, we found that low temperatures enhanced the disease resistance of Arabidopsis against *Pseudomonas syringae* pv. *tomato* (*Pst*) DC3000. This process required Inducer of CBF expression 1 (ICE1), the core transcription factor in cold-signal cascades. ICE1 physically interacted with Non-expresser of *PR* genes 1 (NPR1), the master regulator of the SA signaling pathway. Enrichment of ICE1 on the *PR1* promoter and its ability to transcriptionally activate *PR1* were enhanced by NPR1. Further analyses revealed that cold stress signals cooperate with SA signals to facilitate plant immunity against pathogen attack in an ICE1-dependent manner. Cold treatment promoted interactions of NPR1 and TGA3 with ICE1, and increased the ability of the ICE1–TGA3 complex to transcriptionally activate *PR1*. Together, our results characterize a previously unrecognized role of ICE1 as an indispensable regulatory node linking low temperature activated- and SA-regulated immunity. Discovery of a crucial role of ICE1 in coordinating multiple signals associated with immunity broadens our understanding of plant–pathogen interactions.

## INTRODUCTION

In nature, plants can be attacked by a wide range of pathogens, including bacteria, fungi, oomycetes, viruses, and nematodes. In response, plants have developed a sophisticated innate immune system during co-evolution. The first layer of the defense system is governed by the plasma membrane (PM)-localized pattern recognition receptors (PRRs). The PRRs recognize conserved pathogen-/damage-/microbe-/herbivore-associated molecular patterns (PAMPs/DAMPs/MAMPs/HAMPs) and trigger the first layer of defense responses–this is known as PAMP-triggered immunity (PTI) (Jones and Dangl, 2006; Dodds and Rathjen, 2010; Couto and Zipfel, 2016; Zhou and Zhang, 2020). PTI is generally believed to be sufficient to prevent non-pathogenic microbes from proliferating *in planta*. However, some successful pathogens can deliver effector proteins to evade or suppress host PTI, allowing them to multiply aggressively (Kinya Nomur and He, 2005; Nomura et al., 2005; Nomura et al., 2006; Boller and Felix, 2009; Macho and Zipfel, 2015; Lopez et al., 2019; Wang et al., 2019; Chen et al., 2021; Wang et al., 2022; Wu and Derevnina, 2023). To counter pathogen inhibition of PTI, plants have evolved intracellular nucleotide-binding leucine-rich repeat receptors (NLRs) that detect cellular effectors, directly or indirectly, and cause strong and robust immune responses. These responses, known as effector-triggered immunity (ETI), can stop virulent pathogens from causing severe disease symptoms (Jones and Dangl, 2006; Dodds and Rathjen, 2010; Deslandes and Rivas, 2012; Ma et al., 2020; Bi et al., 2021; Ngou et al., 2021; Yuan et al., 2021; Ngou et al., 2022). Along with the two branches of immune system, the defense hormone salicylic acid (SA) plays a central role in these responses (Spoel and Dong, 2012; Rekhter et al., 2019; Torrens-Spence et al., 2019; Wang et al., 2020; Zavaliev et al., 2020; Peng et al., 2021; Kim et al., 2022; Kumar et al., 2022b; Jia et al., 2023). For example, the accumulation of SA at infection sites subsequently leads to systemic acquired resistance (SAR) against *Pseudomonas syringae* throughout the plant to limit pathogen propagation (Durrant and Dong, 2004; Vlot et al., 2009; Bernsdorff et al., 2016; Hartmann and Zeier, 2019; Kachroo et al., 2020; Zeier, 2021).

In the SA signaling transduction pathway, NONEXPRESSOR OF PATHOGENESIS-RELATED GENES 1 (NPR1) has been well documented as one of the SA receptors that perceives and transduces SA signals (Wang et al., 2020; Kumar et al., 2022a). In the resting state, NPR1 exists in the cytoplasm as an oligomer. After induction, NPR1 monomers are released from oligomers and translocated to the nucleus, where they act as cofactors of transcription factors, such as the TGACG motif-binding transcription factors (TGAs), which regulate global transcriptional reprogramming and resistance to a broad spectrum of pathogens (Zhang et al., 1999; Després et al., 2000; Zhou et al., 2000; Johnson et al., 2003; Kumar et al., 2022a). Recently, NPR1 has attracted attention because of its central roles in plant growth and development, as well as in plant immune responses. For instance, it was reported that NPR1 interferes with the binding of ETHYLENE-INSENSITIVE 3 (EIN3), a key transcription factor involved in ethylene (ET) signaling, to the promoter regions of its target genes to mediate antagonism between SA and ET in apical hook formation (Huang et al., 2020). The transcriptional activation activity of EIN3 is enhanced by NPR1 in a process coordinated by SA and ET during the modulation of leaf senescence (Wang et al., 2021; Yu et al., 2021). Moreover, NPR1 promotes the polyubiquitination and degradation of GID1, the gibberellin receptor, to balance growth and defense (Yu et al., 2022). NPR1 is also involved in facilitating cell survival during ETI by ubiquitinating key proteins involved in immune responses, such as ENHANCED DISEASE SUSCEPTIBILITY 1 and WRKY transcription factors (Zavaliev et al., 2020). NPR1 inhibits the transcriptional activation activity of MYC2, the master regulator of jasmonic acid signaling pathway, to suppress pathogen virulence (Nomoto et al., 2021).

The TGA transcription factors are members of the basic leucine zipper (bZIP) family of proteins, which specifically bind to the TGACG-core activation sequence 1 (*as-1*) to regulate the transcriptional levels of their target genes (Després et al., 2000; Zhou et al., 2000; Johnson et al., 2003). Ten TGA transcription factors are encoded in the Arabidopsis genome, of which TGA1 to TGA7 have been characterized in terms of their ability to interact with NPR1 (Zhang et al., 1999; Després et al., 2000). Among them, TGA3 is mainly required for basal resistance and *PR* gene expression (Kesarwani et al., 2007; Kumar et al., 2022a), and also integrates with other signals. For instance, TGA3 interacts with ARR2, the cytokinin-activated transcription factor, to promote plant immunity through NPR1/TGA3-dependent SA signaling (Choi et al., 2010). A protein complex of TGA3 and WRKY53 acts on the *Cestrum yellow leaf curling virus* promoter to achieve synergistic up-regulation of SA- associated gene expression (Sarkar et al., 2018). BRASSINOSTEROID INSENSITIVE2 (BIN2) balances SA-induced immunity and brassinosteroid-mediated plant growth through phosphorylating TGA3 to activate gene expression (Han et al., 2022).

Low temperature has significant effects on plant growth and development (Orvar et al., 2000; Wang et al., 2009). The ICE1/C-REPEAT BINDING FACTOR/DRE-BINDING FACTOR 1 (ICE1/CBF/DREB1) pathway plays an essential role in plants’ tolerance to low temperature stress, and ICE1 acts as an important transcriptional regulator of cold-responsive gene expression and cold tolerance (Stockinger et al., 1997; Kim et al., 2015). ICE1, a basic helix-loop-helix (bHLH) protein in the MYC class, directly binds to the MYC site (CANNTG) in the promoter of *CBF3* to induce its expression under cold stress (Gilmour et al., 1998; Chinnusamy et al., 2003). ICE1 also binds to the promoters of other genes encoding regulatory factors such as cold-responsive (COR), calmodulin binding transcription activator 2, mitogen-activated protein kinase 3 (MPK3), MPK4, and high expression of osmotically responsive genes 1 (HOS1), all of which are critical for the cold response (Tang et al., 2020). As an essential component of the cold signaling network, ICE1 functions as a key cross-talk node between low temperature signals and other signals. For instance, HOS1, SUMO E3 ligase (SIZ1), Open Stomata 1 (OST1), MPK3/6, and BIN2 are involved in regulating responses to freezing by maintaining homeostasis of the ICE1 protein under cold stress (Dong et al., 2006; Miura et al., 2007; Ding et al., 2015; Li et al., 2017; Zhao et al., 2017; Ye et al., 2019). Moreover, ICE1 integrates endogenous and environmental signals to mediate diverse physiological processes, such as seed germination, flowering time, and stomatal development (Lee et al., 2015, 2017; Hu et al., 2019; An et al., 2021; An et al., 2022).

Plant–pathogen interactions are affected by the ambient temperature, and low temperature can promote plant immune responses. For example, under cold temperatures, winter rye (*Secale cereale*) accumulates abundant pathogenesis-related (PR) proteins that help protect against pathogen attack (Griffith and Yaish, 2004). Singh (2014) found that 7 days of repeated cold stress activated the PTI immune response of Arabidopsis in a histone acetyltransferase 1 (HAT1)-dependent manner, thereby enhancing resistance to the virulent pathogen *Pseudomonas syringae* pv. *tomato* (*Pst*) DC3000. Wu (2019) reported that low temperatures induced a transient defense response to *Pst* DC3000 in plants. Specifically, low temperatures induced H_2_O_2_ accumulation, callose deposition, and defense gene expression (Wu et al., 2019). Notably, there is a growing body of evidence showing that SA-mediated defense responses are activated at low temperature in the interaction between Arabidopsis and *Pseudomonas syringae*. In particular, SA biosynthetic genes, including *ISOCHORISMATE SYNTHASE1* (*ICS1*), *ISOCHORISMATE SYNTHASE2* (*ICS2*), *CALMODULIN BINDING PROTEIN 60-LIKE.g* (*CBP60g*), *SAR DEFICIENT1* (*SARD1*), and *PHENYLALANINE AMMONIA LYASE1* (*PAL1*), are up-regulated by cold stress, activating the SA-mediated immune pathway (Kim et al., 2013; Kim et al., 2017; Wu et al., 2019; Li et al., 2020). Cold temperatures induce expression of the SA signaling-responsive genes *PR1*, *PR2,* and *PR5,* whose encoded products increase plant immunity (Seo et al., 2010). These results suggest that there are broad signaling interactions between low temperature signals and SA responses. Nonetheless, the underlying molecular mechanisms and signaling pathways are unknown.

In this study, we found that the enhancement of plant immune responses by low temperatures largely depends on ICE1, the core transcription factor in the cold signaling pathway. Mechanistic analyses revealed that the ICE1 protein physically interacts with the SA receptor NPR1 and is involved in SA-mediated resistance against *Pst* DC3000. Further analyses demonstrated that ICE1 directly activates the transcription of *PR1* and NPR1 promotes its activities. We also found that ICE1 interacts with TGA transcription factors to cooperatively stimulate *PR1* expression under low temperatures. Taken together, our results demonstrate that the NPR1-TGA3/ICE1 regulatory module acts as a crucial node integrating cold signals and SA signaling in the synergistic regulation of plant immunity.

## RESULTS

### ICE1 Is Involved in Low Temperature-enhanced Plant Immunity

Low temperature has complicated effects on plant immunity (Yang and Hua, 2004; Huang et al., 2010; Yang et al., 2010; Wigge, 2013). To investigate the underlying relationship between low temperature and plant immunity, we assessed effect of low temperature on plant immune responses against the hemibiotropic pathogenic bacteria, *Pseudomonas syringae* pv. *tomato* (*Pst*) DC3000. For these analyses, we monitored disease symptoms and populations of *Pst* DC3000 in Arabidopsis Columbia-0 (Col-0) plants that were grown at normal temperature (22°C) for 4 weeks and then exposed to a long-term (3 weeks) or short-term (10 hours) cold treatment (4°C) before infiltration with the pathogen. We observed that the pathogen-related water-soaking symptom, one of the earliest and most common symptoms of phyllosphere bacterial diseases, as well as bacterial proliferation in infected leaves, were largely reduced in Col-0 plants exposed to a short or long cold treatment, compared with those kept at 22°C (Fig. 1 A–D). These results confirmed that both long- and short-term low temperature treatments promote plant immunity against *Pst* DC3000, consistent with the results of previous studies (Seo et al., 2010; Kim et al., 2017; Wu et al., 2019).

**Figure 1.**
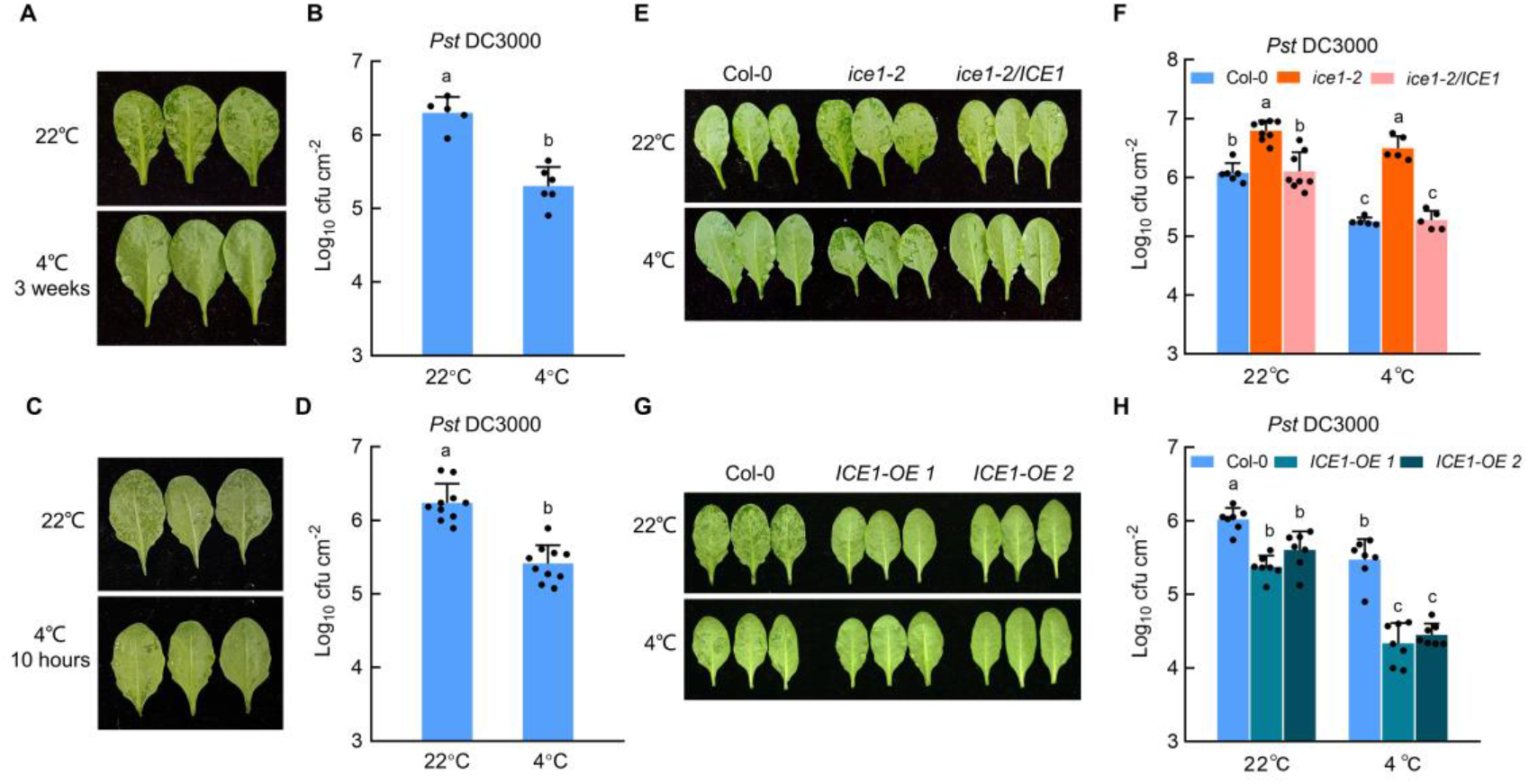
ICE1 involves in low temperature-enhanced plant immunity. (A) Col-0 leaves infiltrated with *Pst* DC3000 (OD_600_ = 0.0001) after 3 weeks of 4°C treatments (down), 22°C as control (up). (B) Bacterial populations in leaves described in (A). Values are displayed as mean ± s.d. (*n* ≥ 5 biological replicates). (C) Col-0 leaves infiltrated with *Pst* DC3000 (OD600 = 0.0001) after 10 hours of 4°C treatments (down), 22°C as control (up). (D) Bacterial populations on leaves described in (C). Values are displayed as mean ± s.d. (*n* = 10 biological replicates). (E) Col-0, *ice1-2*, *ice1-2/ICE1* leaves infiltrated with *Pst* DC3000 (OD_600_ = 0.0001) after 10 hours of 4°C treatments (down), 22°C used as control (up). (F) Bacterial populations on leaves described in (E). Values are displayed as mean ± s.d. (*n* ≥ 5 biological replicates). (G) Col-0, *ICE1-OE 1* and *ICE1-OE 2* leaves infiltrated with *Pst* DC3000 (OD_600_ = 0.0001) after 10 hours of 4°C treatment (down), 22°C as control (up). (H) Bacterial populations of *Pst* DC3000 on plants described in (G). Values are displayed as mean ± s.d. (*n* = 7 biological replicates). Photos were taken after 48 h pathogen inoculation. Different letters indicate statistically significant differences (two-way ANOVA, *P* < 0.05). Experiments were repeated at least three times with similar trends.

ICE1 is a transcriptional inducer of genes that encode important components of chilling tolerance of Arabidopsis (Chinnusamy et al., 2003; Lee et al., 2005; Benedict et al., 2006; Chinnusamy et al., 2007). At the early stages of cold stress, ICE1 in its phosphorylated form activates the expression of *CBFs* to enhance plant freezing tolerance (Ding et al., 2015; Ye et al., 2019). Given the critical role of ICE1 at the early stage of the cold stress response, we hypothesized that ICE1 may be involved in short-term low temperature-enhanced immunity. To test this possibility, we investigated *Pst* proliferation in Col-0, the loss-of-function *ice1-2* mutant, and an *ICE1*-complemented mutant (*ice1-2/ICE1*) after short-term low temperature treatment. As shown in Fig. 1E, F, the leaves of Col-0 and *ice1-2/ICE1* exhibited clearly enhanced disease-resistance phenotypes, with less severe water-soaking symptoms and lower pathogen populations after cold treatment, compared with the control. However, in *ice1-2* mutants, the water-soaking symptoms were severe and the pathogen population was large (Fig. 1E, F). These results suggest that ICE1 may positively regulate cold stress-enhanced plant immunity. To confirm the role of ICE1, we used two stable *ICE1*-overexpressing transgenic lines, *35S::GFP-ICE1* (Chinnusamy et al., 2003); hereinafter *ICE1-OE 1*) and *35S::3Myc-ICE1* (hereinafter *ICE1-OE 2*) for bacterial propagation assays. As expected, compared with Col-0, the *ICE1*-overexpressing plants showed enhanced resistance, including less severe water soaking symptoms and lower pathogen proliferation after low temperature stress (Fig. 1G, H). Together, these results indicate that ICE1 acts as a positive regulator of plant immunity under ambient and cold temperatures.

### ICE1 Physically Interacts with NPR1

ICE1 regulates diverse physiological processes through associating with other proteins (Ding et al., 2015; Lee et al., 2015; Hu et al., 2019; Ye et al., 2019; An et al., 2021; An et al., 2022). To explore the underlying molecular mechanism by which ICE1 regulates plant immune responses, we performed yeast two-hybrid screening from a pool of key regulatory proteins that act in Arabidopsis immunity, such as proteins that are involving in SA signaling pathway and PTI response. From this screening, a potential candidate, NPR1, displayed strong interaction with ICE1 in the yeast cells and selected as the first potential candidate for further characterization (Fig. 2A). To identify the functional region responsible for the interaction between ICE1 and NPR1, two truncated versions of the proteins were used in yeast two-hybrid assays. As shown in Fig. 2A, deletion of the N-terminal amino acid residues 1–260 of ICE1 (AD-ICE1^261-249^) did not affect the ICE1–NPR1 interaction, whereas deletion of the C-terminal residues of ICE1 containing the bHLH domain (AD-ICE1^1-260^) eliminated the interaction completely (Fig. 2A). Additionally, the N-terminal amino acid residues of NPR1 containing the BTB structural domain (BD-NPR1^1-194^) strongly interacted with ICE1, while the C-terminal fragment (BD-NPR1^178-593^) did not. These results demonstrate that the bHLH structural domain of ICE1 and the BTB structural domain of NPR1 are necessary for the ICE1–NPR1 interaction.

**Figure 2.**
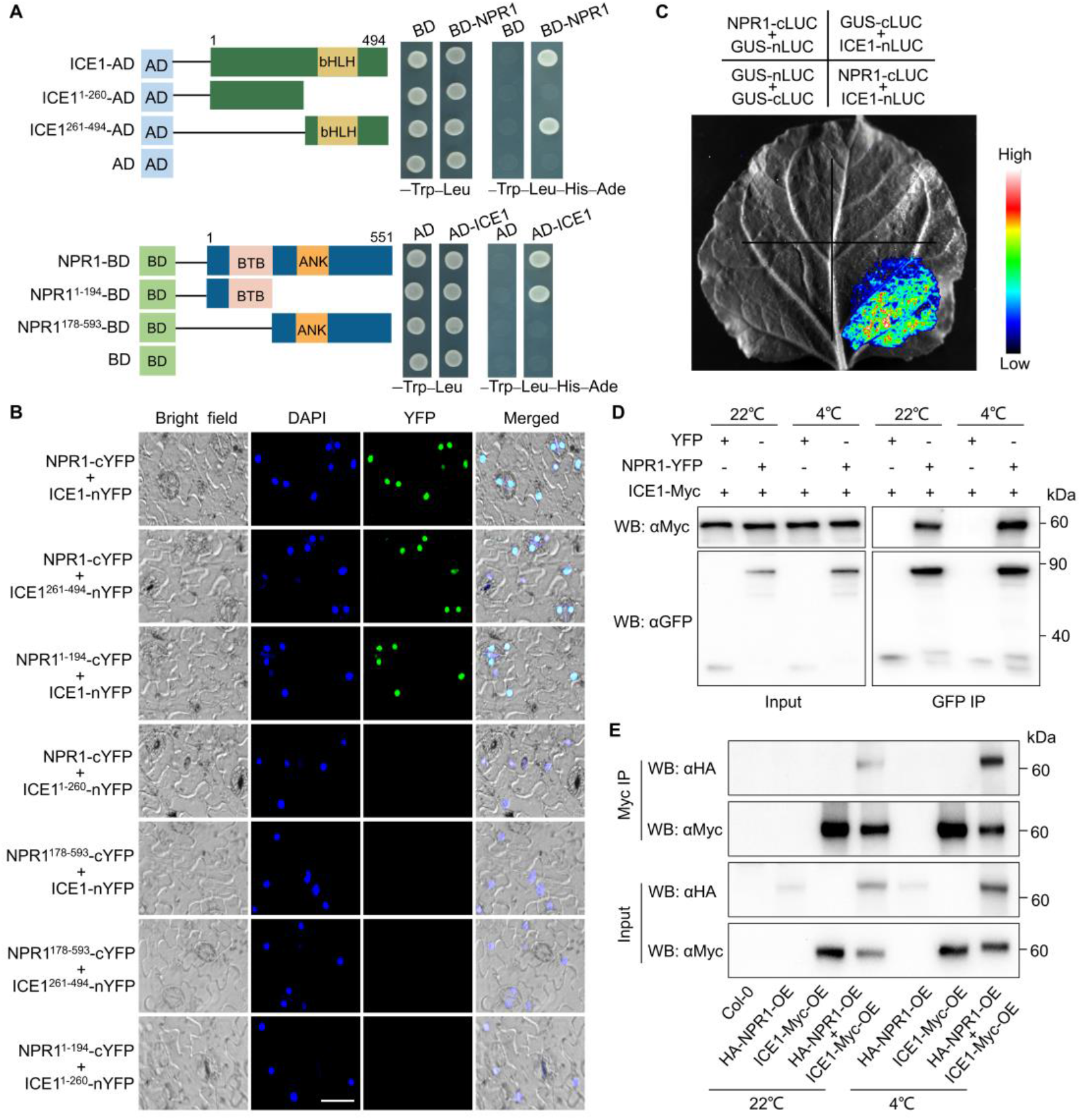
ICE1 interacts with NPR1. (A) Yeast two hybrid assays showing the interactions between ICE1, NPR1, and their truncated versions. Protein interactions were indicated by the growth of yeast cells after 2 days of incubation in dropout medium lacking Leu, Trp, His and Ade. The numbers indicate the positions of amino acids, pGBKT7 (BD) and pGADT7 (AD) were used as negative controls. (B) BiFC assay. Fluorescence was observed in the nucleus of transformed *N. benthamiana* cells co-expressing ICE1-nYFP (or ICE1^261-494^-nYFP) with NPR1-cYFP or ICE1-nYFP with NPR1^1-194^-cYFP at 4°C for 4 h. No signal was obtained in the negative controls which NPR1-cYFP (or NPR1^1-194^-cYFP) with ICE1^1-260^-nYFP or ICE1-nYFP (or ICE1^261-494^-nYFP) with NPR1^178-593^-cYFP were co-expressed. Nuclei are indicated by DAPI staining. Scale bar = 20 μm. (C) Interaction between ICE1 and NPR1 in split luciferase assay. ICE1-nLUC and NPR1-cLUC were co-expressed in *N. benthamiana* leaves, the luminescence intensity was detected at 4°C for 4 h after 48 h of incubation by an imaging system. GUS-nLUC and GUS-cLUC were set as a negative control. (D) Co-IP assay. Arabidopsis protoplasts expressing NPR1-YFP with ICE1-Myc, YFP with ICE1-Myc were incubated for 16 h at 22°C. For low-temperature induction, protoplasts were processed at 4°C for 2 h before total protein was extracted. Then total protein was immunoprecipitated with GFP-trap agarose and the co-immunoprecipitated proteins were detected with anti-Myc antibody. Experiments were repeated at least three times with similar trends. (E) Co-IP assay using stable transgenic seedlings. Transgenic plants are growing on a normal temperature chamber for 4 weeks, Col-0 plants used as negative control. For low temperature treatment, plants are moved into another growth chamber with 4°C for 2 h treatment before protein extraction. ICE1-Myc fusion was immunoprecipitated by Myc-trap agarose, and the coimmunoprecipitated protein was detected using an anti-HA antibody. Experiments were repeated at least three times with similar trends.

Bimolecular fluorescence complementation (BiFC) assays were conducted to further confirm the interaction between ICE1 and NPR1. For these assays, NPR1 fused to the C-terminal of YFP (NPR1-cYFP) and ICE1 fused to the N-terminal of YFP (ICE1-nYFP) were co-expressed in *Nicotiana benthamiana* leaves, and then fluorescence in the leaves was observed under a laser confocal microscope. Almost no YFP fluorescence was observed when NPR1-cYFP was co-expressed with ICE1-nYFP in *N. benthamiana* leaves at 22°C (Supplemental Fig. S1A), but the fluorescence signal became very strong after induction at 4°C for 4 h (Fig. 2B; Supplemental Fig. S2A), suggesting that the association between NPR1 and ICE1 may depend on low temperature *in vivo*. To further verify this interaction, we performed a split luciferase assay by fusing these proteins to the C-terminus (cLUC) or N-terminal (nLUC) of luciferase, with LUC fused with β-glucuronidase (GUS) as the negative control. Similarly, very faint LUC activity was detected when NPR1-cLUC was transiently co-expressed with ICE1-nLUC in *N. benthamiana*, but strong LUC signals were observed after low temperature induction (Fig. 2C; Supplemental Fig. S1B).

Consistently, in a co-immunoprecipitation (Co-IP) assay, ICE1 was immunoprecipitated by anti-GFP agarose beads from Arabidopsis protoplasts co-expressing NPR1-YFP and ICE1-Myc, but not from those co-expressing YFP and ICE1-Myc, and more precipitation products were detected after low temperature induction (Fig. 2D). To further confirm the interaction between ICE1 and NPR1 *in vivo*, we performed Co-IP assay using stable Arabidopsis transgenic lines both under normal and cold temperature conditions. In line with the results conducted in protoplasts, more NPR1 proteins could be co-immunoprecipitated with ICE1 after cold temperature treatment (Fig. 2E). Taken together, these results indicate that ICE1 interacts with NPR1 both *in vitro* and *in vivo*, and that low temperature potentiates the association between ICE1 and NPR1 *in planta*.

### ICE1 Is Required for SA-mediated Plant Immunity

Because ICE1 physically associates with the SA receptor NPR1 (Fig. 2), we hypothesized that ICE1 may participate in SA signal transduction. To test this hypothesis, we first detected changes in the transcript levels of *ICE1* in response to benzothiadiazole (BTH), a synthetic analog of SA that effectively induces SA responses and is extensively used as an alternative to SA signaling (Huot et al., 2017; Kim et al., 2022). Interestingly, BTH treatment did not affect the transcript level of *ICE1*, but led to the accumulation of the ICE1 protein to high levels (Supplemental Fig. S3). Next, we examined whether ICE1 is involved in SA-regulated immunity by conducting a BTH protection assay. For this assay, wild-type Col-0, *ice1-2,* and *ice1-2/ICE1* plants were pre-treated with BTH for 24 h before pathogen infiltration, and then their susceptibility to disease caused by *Pst* DC3000 was evaluated. Compared with the mock pretreated plants, the BTH-pretreated Col-0 and *ice1-2/ICE1* plants exhibited greatly reduced disease symptoms on the leaves after *Pst* DC3000 infection (Fig. 3A and 3B). In contrast, the leaves of *ice1-2* displayed severe disease symptoms, i.e., severe water soaking symptoms and strong pathogen growth, after *Pst* DC3000 infection (Fig. 3A, B). The transcript levels of *PR1*, *PR2*, *PR5* and *WRKY70* were also lower in BTH-pretreated *ice1-2* plants (Fig. 3C and Supplemental Fig. S4). These results suggest that ICE1 may be required for effective SA-regulated immunity. To further confirm the essential role of ICE1 in SA-regulated immunity, we analyzed the disease symptoms of pretreated Col-0 and *ICE1*-overexpressing plants (*ICE1-OE 1* and *ICE1-OE 2*). Compared with Col-0 plants, BTH-pretreated *ICE1*-overexpressing plants showed less severe disease symptoms after inoculation with *Pst* DC3000, accompanied by BTH-induced high transcript levels of *PR1* (Fig. 3D–F).

**Figure 3.**
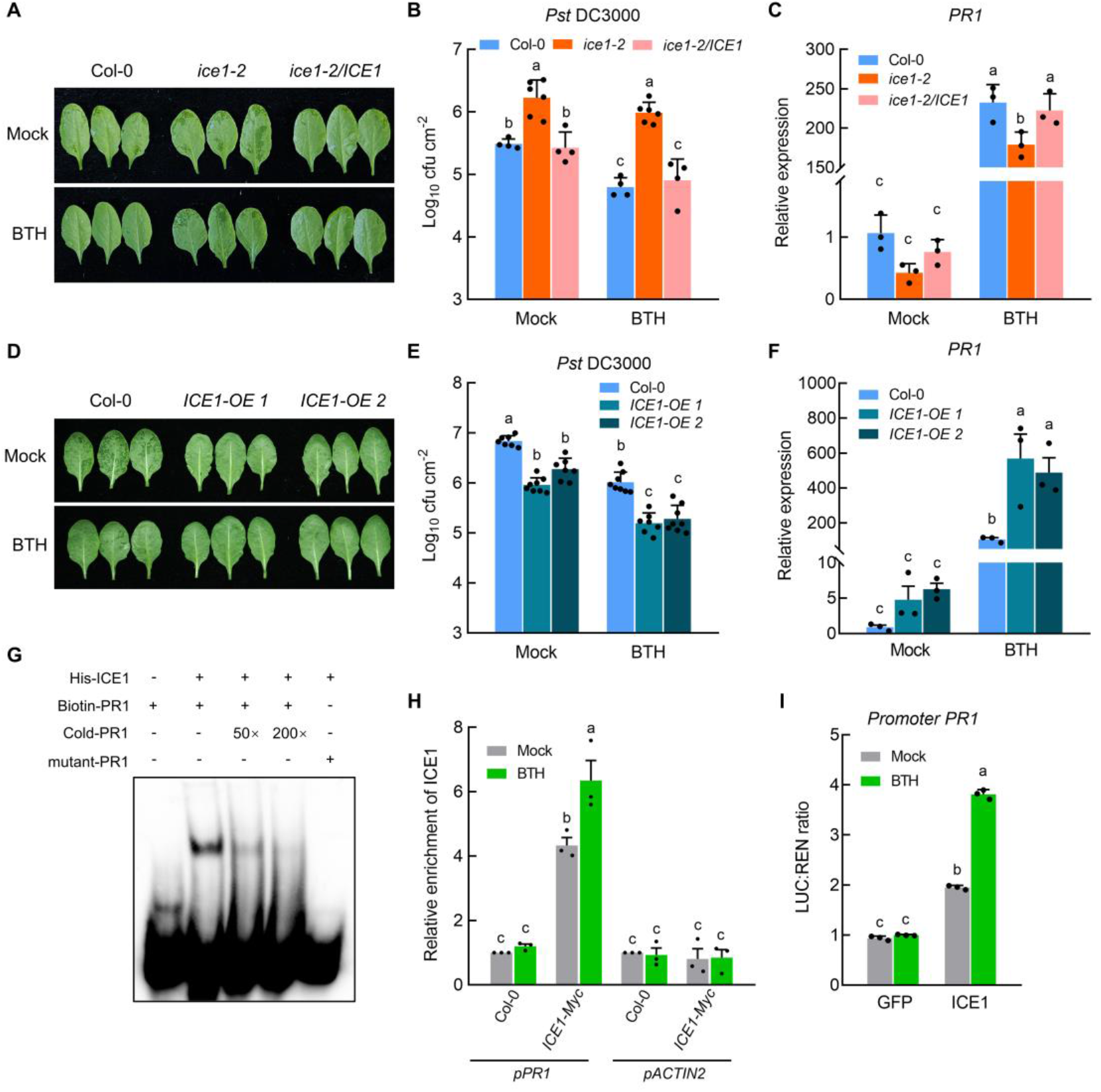
ICE1 involves in SA-regulated immunity. (A) Col-0, *ice1-2*, *ice1-2/ICE1* leaves infiltrated with *Pst* DC3000 (OD_600_ = 0.0001) after 24 hours pretreating with 100 μM BTH (down), mock as control (up). (B) Bacterial populations in leaves described in (A). Values are displayed as mean ± s.d. (*n* ≥ 4 biological replicates). (C) RT-qPCR analysis of *PR1* expression in leaves described in (A). Values are displayed as mean ± s.e.m. (*n* = 3 biological replicates). The *ACTIN2* gene was used as a control. (D) Col-0, *ICE1-OE 1* and *ICE1-OE 2* leaves infiltrated with *Pst* DC3000 (OD_600_ = 0.0001) after 24 hours pretreating with mock (up) and 100 μM BTH (down). (E) Bacterial populations in leaves described in (D). Values are displayed as mean ± s.d. (*n* ≥ 7 biological replicates). (F) RT-qPCR analysis of *PR1* expression in leaves described in (D). Values are displayed as mean ± s.e.m. (*n* = 3 biological replicates). The *ACTIN2* gene was used as a control. (G) EMSA assay examines ability of ICE1 binding to the E-box (CANNTG) of *PR1* promoter. Biotin-labeled probes were incubated with ICE1-His proteins, and the free and bound DNAs were separated on an acrylamide gel. As indicated, unlabeled probe (cold-*PR1*) was used as competitor and biotin labeled mutant probe (mutant-*PR1*) was used as negative control. Mutated probe in which the CATTTG motif was replaced with AATTTC. (H) ChIP-qPCR analysis of the relative enrichment of ICE1 on the promoter of *PR1*. Col-0 and *ICE1*-overexpressing (*ICE1-Myc*) plants were treated with mock or 100 μM BTH for 6 h and pooled for ChIP assays using anti-Myc antibody, the *ACTIN2* untranslated region sequence (*pACTIN2*) as a negative control. Values are displayed as mean ± s.e.m. (*n* ≥ 3 biological replicates). (I) Transient dual-LUC reporter assays showing that ICE1 activates the expression of *PR1*. Values are displayed as mean ± s.e.m. (*n* ≥ 3 biological replicates). Each biological replicate was from different leaves of more than 60 Col-0 plants. Photos were taken after 48 h pathogen inoculation in A and D. Different letters indicate statistically significant differences (two-way ANOVA, *P* < 0.05). Experiments were repeated at least three times with similar trends.

*PR1*, whose expression is activated by the NPR1–TGA regulation module, is a principal output gene of SA-associated immunity (Zhang et al., 1999; Després et al., 2000; Zhou et al., 2000; Johnson et al., 2003). Given that ICE1 interacts with NPR1 and plays an essential role in the SA signaling pathway (Figures 2 and 3), we wondered whether ICE1 directly regulates the transcription of *PR1* in a similar manner as TGA transcription factors. ICE1 is a MYC-like bHLH transcriptional activator that binds specifically to the MYC recognition sequences (CANNTG, also known as the E-box) in the *CBF3* promoter (Chinnusamy et al., 2003; Tang et al., 2020). We analyzed the promoter of *PR1* and found that some MYC recognition sequences exist in the promoter of *PR1*, 1000 bp upstream of ATG (Supplemental Fig. 5). Hence, we performed electrophoresis mobility shift assays (EMSAs) to determine whether ICE1 could bind directly to the E-box *cis*-element in the promoter of *PR1*. As shown in Fig. 3G, the ICE1 protein was able to bind to the E-box in the *PR1* promoter (Fig. 3G). To verify these findings, we conducted chromatin immunoprecipitation (ChIP) assays with *ICE1-Myc* (*ICE1-OE 2*) plants treated or not with BTH. We observed that ICE1 was enriched in the *PR1* promoter region, and BTH promoted its enrichment (Fig. 3H). Next, a dual-luciferase reporter assay was conducted to determine the effects of ICE1 on *PR1* transcription. *LUC* expression driven by the *PR1* promoter was significantly activated by ICE1 (Fig. 3I). Taken together, these results indicate that ICE1 is required for effective SA-associated immunity *via* direct activation of *PR1* transcription.

### NPR1 Enhances the Ability of ICE1 to Transcriptionally Activate *PR1*

Having established that ICE1 interacts with NPR1 and positively regulates SA responses through direct binding to the promoter region of *PR1*, we investigated whether ICE1 functions in a NPR1-relevant manner. To test this possibility, we generated *ice1-2 npr1-1* double mutant plants by crossing the *ice1-2* mutant with *npr1-1*, a loss-of-function mutant allele of *NPR1* (Col-0 background;(Cao et al., 1997)). Similar to the *ice1-2* and *npr1-1* single mutants, the *ice1-2 npr1-1* double mutant displayed a highly susceptible phenotype to *Pst* DC3000 (Fig. 4A, B), implying that ICE1 may act in the same pathway as NPR1 to mediate plant immunity.

**Figure 4.**
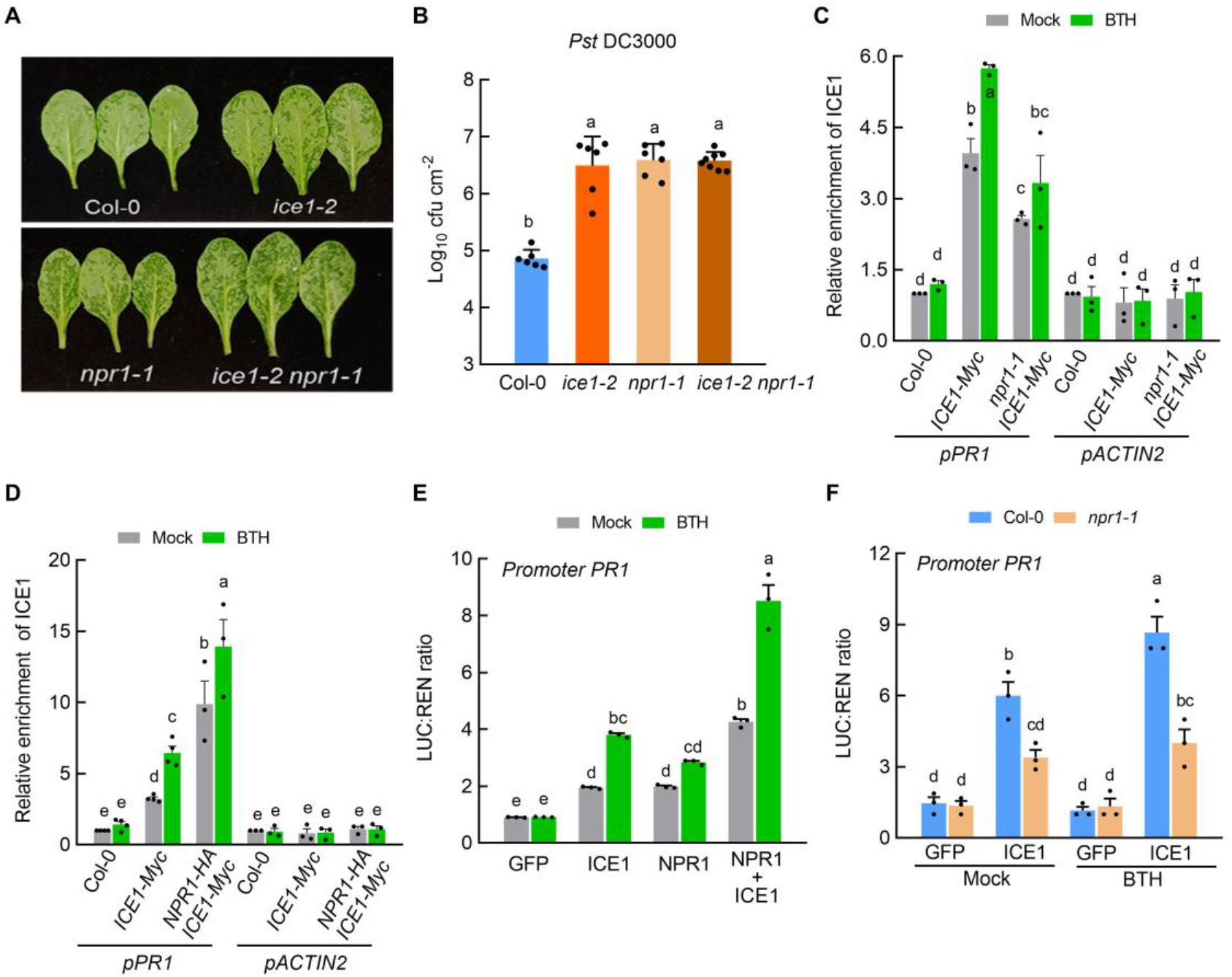
NPR1 promotes the binding and transcriptional activation of ICE1 to *PR1*. (A) Col-0, *ice1-2*, *npr1-1* and *ice1-2 npr1-1* leaves infiltrated with *Pst* DC3000 (OD_600_ = 0.0001). Photos were taken after 48 h pathogen inoculation. (B) Bacterial populations in leaves described in (A). Values are displayed as mean ± s.d. (*n* ≥ 6 biological replicates). (C and D) ChIP-qPCR analysis of the relative enrichment of ICE1 on the promoter regions of *PR1*. Col-0, *ICE1*-overexpressing (*ICE1-Myc*), *npr1-1 ICE1-Myc* and *NPR1-HA ICE1-Myc* plants were treated with mock or 100 μM BTH for 6 h and pooled for ChIP assays using anti-Myc antibody, the *ACTIN2* untranslated region sequence (*pACTIN2*) as a negative control. Values are displayed as mean ± s.e.m. (*n* = 3 biological replicates). (E) Transient transcriptional activity assays showing that NPR1 enhances the transcriptional activation of ICE1 to *PR1*. Values are displayed as mean ± s.e.m. (*n* = 3 biological replicates). (F) Transient transcriptional activity assays showing that activation of the *PR1* promoter by ICE1 is compromised in the *npr1-1* mutant. Values are displayed as mean ± s.e.m. (*n* = 3 biological replicates). Each biological replicate was from different leaves of more than 60 plants in (E) and (F). Different letters indicate statistically significant differences (two-way ANOVA, *P* < 0.05). Experiments were repeated at least three times with similar results.

NPR1 has long been believed to associate with downstream transcription factors, such as TGAs, to trigger their transcriptional activation of *PR* genes (Zhang et al., 1999; Després et al., 2000; Zhou et al., 2000; Johnson et al., 2003; Kumar et al., 2022a). Thus, we wondered whether NPR1 could promote the ability of ICE1 to activate the transcription of *PR1*. To test this possibility, ChIP assays were conducted with *npr1-1 ICE1-Myc* and *NPR1-HA ICE1-Myc* plants treated or not with BTH. As shown in Fig. 4C, D, ICE1 was enriched on the *PR1* promoter in *ICE1-Myc*, and BTH promoted its enrichment. However, the high enrichment of ICE1 on the promoter of *PR1* was impaired in the background of *npr1-1*, but enhanced in the *NPR1*-overexpressing line (Fig. 4C, D). These results suggest that the enrichment of ICE1 on the promoter of *PR1* is promoted by NPR1.

Next, dual-luciferase reporter assays using Arabidopsis mesophyll protoplasts were conducted to validate the effect of NPR1 on the transcriptional activation activity of ICE1. The effectors consisted of GFP, NPR1, and ICE1 under the control of *Pro35S*, and the reporter consisted of the *PR1* promoter fused to the *LUC* gene (Supplemental Fig. S6). Interestingly, *LUC* expression driven by the *PR1* promoter was also induced when NPR1 and GFP were co-expressed, implying that basal TGA or other proteins were involved. However, *LUC* expression was induced to a much higher level when NPR1 and ICE1 were co-expressed than when GFP and ICE1 were co-expressed (Fig. 4E). In contrast, *LUC* expression driven by the *PR1* promoter was compromised by the loss-of-function of *NPR1* (Fig. 4F). These results provide further evidence that the ability of ICE1 to activate the transcription of *PR1* is stimulated by NPR1.

### ICE1 Integrates Low Temperature-enhanced and SA-regulated Immunity

ICE1 is involved in low temperature-activated plant immunity (Fig. 1) and plays a crucial role in the SA signaling pathway (Fig. 3), implying that it may function as a central node linking these two signaling pathways. To test this hypothesis, we assessed disease symptoms, including water-soaking symptoms and pathogen proliferation, in Col-0, *ice1-2,* and *ice1-2*/*ICE1* plants treated with BTH for 24 hours with or without a 10-hour low temperature pretreatment. This allowed us to characterize the critical role of ICE1 in integrating low temperature signals and SA signals to induce pathogen resistance. As shown in Fig. 5A, B, in the leaves of Col-0 and *ice1-2/ICE1,* the water soaking symptoms and pathogen proliferation were alleviated by either BTH or 4°C pretreatment, and significantly diminished by BTH combined with cold treatment, compared with the mock treatment. In contrast, the leaves of *ice1-2* showed severe water soaking symptoms and high pathogen proliferation under all the different treatments. The transcript levels of *PR1* were also lower in *ice1-2* plants compared with Col-0 and *ice1-2/ICE1* plants under different conditions (Fig. 5C). We tested the enrichment of ICE1 on the promoter of *PR1* in the different treatments. As shown in Fig. 5D, accumulation of ICE1 on the promoter of *PR1* was enhanced by the combined low temperature and BTH treatment, compared with either of these treatments alone. Collectively, these results indicate that ICE1 functions as a crucial node by integrating low temperature signals and SA signals to promote plant disease immunity synergistically.

**Figure 5.**
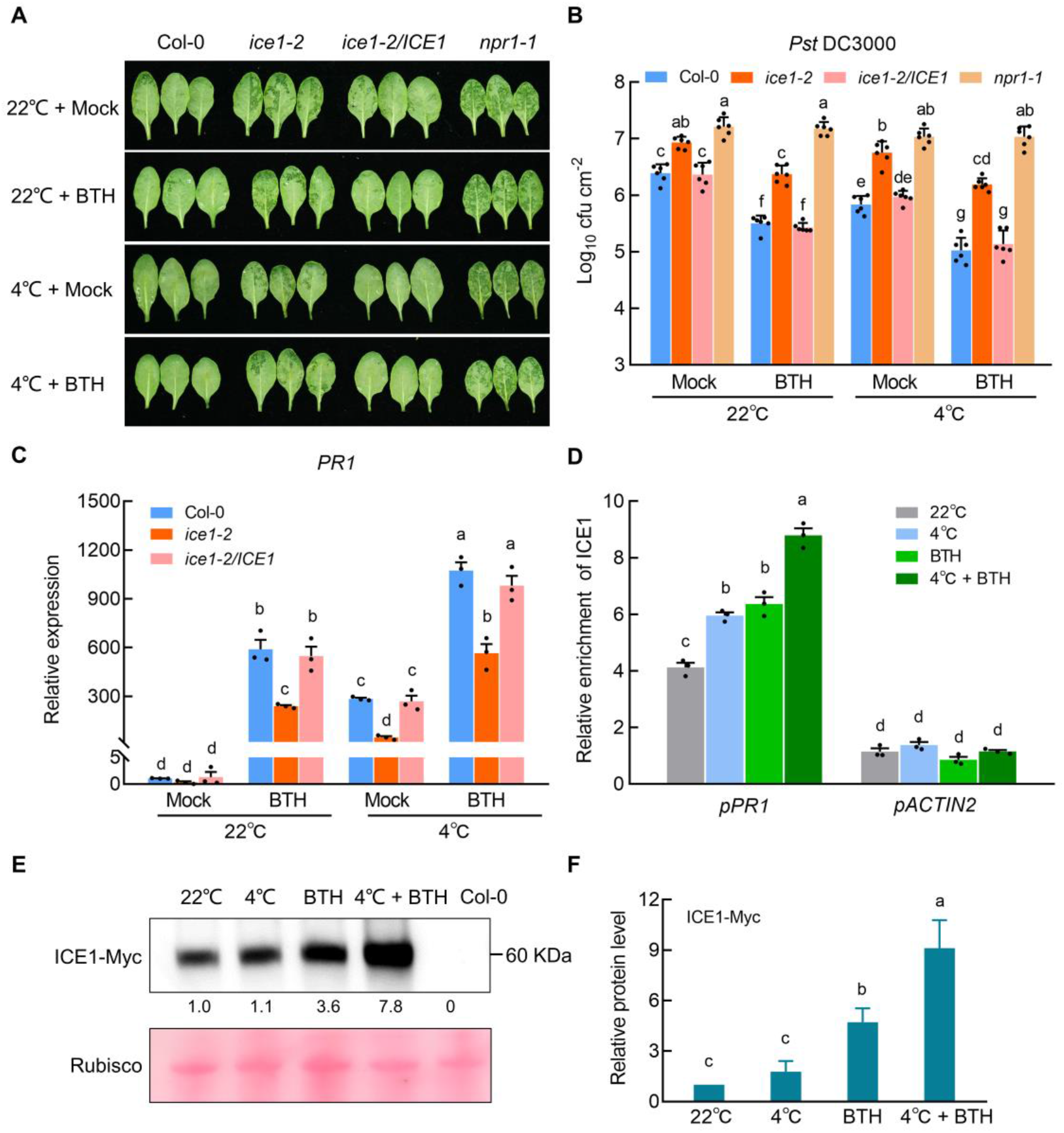
ICE1 links low temperature signals and SA pathway to promote plant immunity. (A) Col-0, *ice1-2* and *ice1-2/ICE1* leaves infiltrated with *Pst* DC3000 (OD_600_ = 0.0001) after BTH combined with low temperature pretreating (briefly, 4-week-old plants were sprayed with mock or BTH, and kept them in a growth chamber with normal temperature (22°C) for 14 h. After then, the low temperature experimental plants were transferred to a low temperature (4°C) growth chamber for 10 h cold treatment.). *npr1-1* mutant plants was used as a control. Photos were taken after 48 h pathogen inoculation. (B) Bacterial populations in leaves described in (A). Values are displayed as mean ± s.d. (*n* = 6 biological replicates). (C) RT-qPCR analysis of *PR1* expression in leaves described in (A). Values are displayed as mean ± s.e.m. (*n* = 3 biological replicates). The *ACTIN2* gene was used as a control. (D) ChIP-qPCR analysis of the relative enrichment of ICE1 on the promoter regions of *PR1*. *ICE1*-overexpressing plants (*ICE1-Myc*) were treated with 100 μM BTH for 6 hours and 4°C for 2 hours and pooled for ChIP assays using anti-Myc antibody, the *ACTIN2* untranslated region sequence (*pACTIN2*) as a negative control. Values are displayed as mean ± s.e.m. (*n* = 3 biological replicates). (E) Immunoblot analyzing the accumulation of ICE1 protein. Ten-day-old *ICE1*-overexpressing seedlings (*ICE1-Myc*) were treated with 100 μM BTH for 6 h and/or 4°C for 2 hours before protein extraction. The accumulation of ICE1 protein was detected with anti-Myc antibody. (F) Relative protein levels of ICE1-Myc in (E), which were quantified by Image J. Values are displayed as mean ± s.e.m. (*n* = 3 biological replicates). Different letters indicate statistically significant differences (two-way ANOVA, *P* < 0.05). Experiments were repeated at least three times with similar trends.

To further explore the underlying molecular mechanism by which ICE1 is synergistically regulated by these two important cues, we determined the amount of ICE1 protein that accumulated in response to a combined low temperature and BTH treatment. Consistent with previous results, ICE1 protein remained stable within 2 hours of 4°C treatment (Ding et al., 2015). Strikingly, ICE1 accumulated to a much higher level under 6 hours BTH treatment combined with 2 hours of 4°C treatment than in either the low temperature or BTH treatment alone (Fig. 5E-F).

### CBF Pathway Does not Play a Major Role in the Low Temperature- and SA-Regulated Immunity

As *CBF3* is the canonical target of ICE1, we are wondering if this ICE1-mediated immunity involves *CBF3* or not. To address this question, we monitored disease symptoms and populations of *Pst* DC3000 in 4-week-old Col-0 and *cbfs* mutant plants that were treated with or without short-term (10 h) cold temperature (4°C). We observed that the pathogen-related water-soaking symptom as well as bacterial proliferation showed no significant differences between Col-0 and *cbfs* mutant leaves at either normal temperature or cold temperature (Supplemental Fig. S7A, B). Expression of *PR1* showed no obvious differences in Col-0 and *cbfs* mutant plants at both normal and low temperature treatments (Supplemental Fig. S7C). These results suggest that *CBF* genes does not involve in low temperature-enhanced immunity, which is consistent with Li’s results that CBF pathway does not play a major role in mediating low-temperature enhancement of disease resistance to *Pst* DC3000 (Li et al., 2020).

To explore if CBFs involves ICE1-regulated immunity, we conducted the BTH protection assay in wild-type Col-0 and *cbfs* mutant plants. However, no obvious differences occurred between them either in disease symptoms and pathogen proliferation or *PR1* expression, suggesting that *CBF* genes did not mediate SA-regulated immunity (Supplemental Fig. S7D-F).

### ICE1 Physically Interacts with TGA Transcription Factors

TGA transcription factors were confirmed to be interacting proteins of NPR1 in yeast two-hybrid screening assays (Zhang et al., 1999; Zhou et al., 2000), and are believed to be the nuclear targets that bind to the *as-1* element required for SA-induced *PR1* gene expression (Zhang et al., 1999; Zhou et al., 2000; Johnson et al., 2003; Kesarwani et al., 2007; Ding et al., 2018). To explore the regulatory role of ICE1 in SA signaling comprehensively, we investigated whether ICE1 interacts with TGA transcription factors. The full-length coding sequences (CDSs) of *TGA1–TGA7* were fused to pGBK-T7 to construct pGBK-TGAs vectors, and then each one was transformed into the yeast strain with pGAD-ICE1. Interestingly, ICE1 interacted with all tested TGA proteins in yeast (Fig. 6A). Among the seven members of the TGA family in Arabidopsis, TGA3 plays an important role to bind to the *as-1* element of *PR1* to induce its expression (Kesarwani et al., 2007; Choi et al., 2010; Kumar et al., 2022a). Hence, we focused on TGA3 in subsequent experiments.

**Figure 6.**
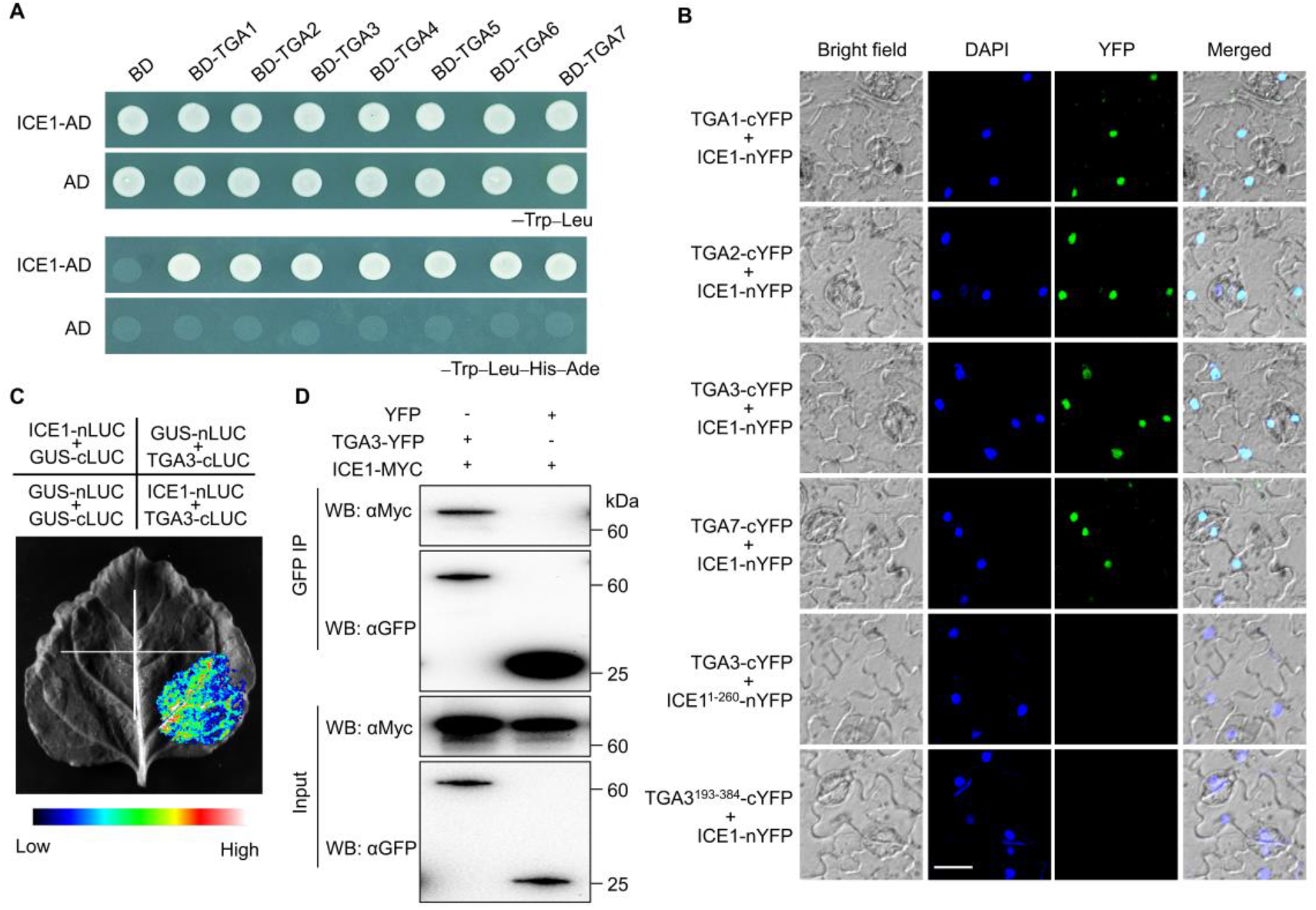
ICE1 interacts with TGA transcription factors. (A) Yeast two hybrid assays. Interactions of ICE1 with TGAs were indicated by the ability of yeast cells to grow on dropout medium lacking Leu, Trp, His, and Ade for 2 days after plating. BD and AD were used as negative controls. (B) BiFC assay. Fluorescence was observed in the nuclear compartment of transformed *N. benthamiana* cells, resulting from the complementation of ICE1-nYFP with TGA1/2/3/7-cYFP. No signal was obtained for the negative controls in which ICE1^1-260^-nYFP with TGA3-cYFP or ICE1-nYFP with TGA3^193-384^-cYFP were coexpressed. Nuclei are indicated by DAPI staining. Scale bar = 20 μm. (C) Split luciferase assay. nLUC-ICE1 and cLUC-TGA3 constructs were co-transformed into *N. benthamiana*. Luminescence intensity was measured after 48 h of incubation by an imaging system. GUS-nLUC and GUS-cLUC were set as a negative control. (D) Co-IP assay. Arabidopsis protoplasts expressing TGA3-YFP with ICE1-Myc, YFP with ICE1-Myc were incubated for 16 h. Total protein was extracted and then immunoprecipitated with GFP-trap agarose. Co-immunoprecipitated protein was detected with anti-Myc antibody. Experiments were repeated at least three times with similar trends.

To identify the functional regions responsible for the interaction between ICE1 and TGAs, different truncated versions of the proteins were used in yeast two-hybrid assays. As shown in Supplemental Fig. S8A, deletion of the N-terminal amino acid residues 1–260 of ICE1 (AD-ICE1^261-494^) did not affect the ICE1–TGA3 interaction, whereas deletion of the C-terminal residues of ICE1 containing the bHLH domain (AD-ICE1^1-260^) eliminated the interaction completely. Additionally, the N-terminal amino acid residues of TGA3 containing the bZIP domain (BD-TGA3-N) strongly interacted with ICE1, while the C-terminal fragment (BD-TGA3-C) did not. These results demonstrate that the bHLH structural domain of ICE1 and the bZIP structural domain of TGA3 are necessary for the ICE1–TGA3 interaction (Supplemental Fig. S8).

The results of the BiFC, split luciferase and Co-IP assays further confirmed that ICE1 interacted with TGA3 in plant cells. For the BiFC assays, full-length CDSs of *TGA1* (clade I), *TGA2* (clade II), *TGA3*, and *TGA7* (clade III) were fused to the C-terminal yellow fluorescent protein (cYFP) fragment to generate TGA1-cYFP, TGA2-cYFP, TGA3-cYFP, and TGA7-cYFP, respectively, and the full-length CDS of *ICE1* was fused to the N-terminal fragment of YFP (nYFP) to produce ICE1-nYFP. Truncated TGA3^193-384^-cYFP and ICE1^1-260^-nYFP that could not interact with ICE1 or TGA3 were used as negative controls (Supplemental Fig. S2). When ICE1-nYFP was transiently co-expressed with TGAs-cYFP in leaf cells of *N. benthamiana*, recombinant YFP fluorescence was observed in the nucleus, whereas no fluorescence signals were detected when either TGA3^193-384^-nYFP and ICE1-nYFP or ICE1^1-260^-nYFP and TGA3-cYFP were co-expressed (Fig. 6B; Supplemental Fig. S2B).

For the split luciferase assay, TGA3 fused to the C-terminus (cLUC) and ICE1 fused to N-terminal (nLUC) of luciferase were co-expressed in *N. benthamiana* leaves. As shown in Fig. 6C, LUC activity could be detected when TGA3 interacted with ICE1. In the Co-IP assay, ICE1 was immunoprecipitated by anti-GFP agarose beads in Arabidopsis protoplasts co-expressing TGA3-YFP and ICE1-Myc, but not in those co-expressing YFP and ICE1-Myc. This result provided further evidence for the interaction between ICE1 and TGA3 *in vivo* (Fig. 6D). Taken together, these results demonstrate that ICE1 physically interacts with the TGA3 transcription factor *in planta*.

### ICE1 and TGA3 Works Synergistically during Low Temperature-enhanced Immunity

Having demonstrated that ICE1 associates with TGA3, we wondered if these two proteins act cooperatively to modulate disease resistance in plants exposed to cold treatment. Hence, we constructed the *TGA3*-knockout mutant *tga3^CR^* (Supplemental Fig. S9), and then crossed *ice1-2* with *tga3^CR^* to obtain *ice1-2 tga3^CR^* double mutant plants. The disease phenotypes of these plants were then determined. As shown in Fig. 7A, B, compared with the *ice1-2* and *tga3^CR^* single mutants, the *ice1-2 tga3^CR^* double mutant showed a more susceptible phenotype, including severe water soaking symptoms and high pathogen proliferation after inoculation with *Pst* DC3000 both at normal and low temperatures, indicating a synergistic role of TGA3 and ICE1 in the regulation of plant immunity.

**Figure 7.**
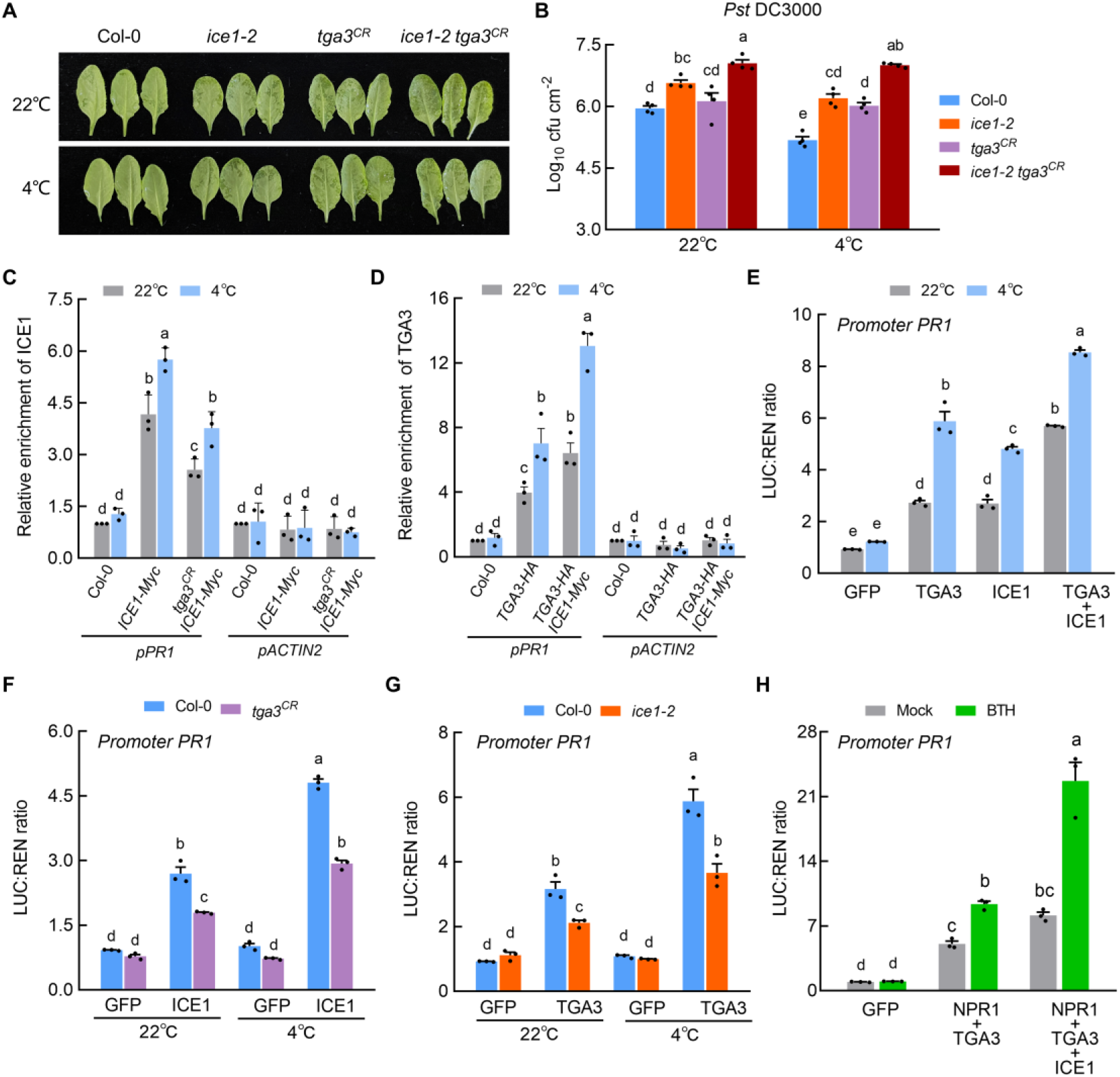
ICE1 and TGA3 works synergistically to activate expression of *PR1* during low temperature stress. (A) Col-0, *ice1-2*, *tga3^CR^* and *ice1-2 tga3^CR^* leaves infiltrated with *Pst* DC3000 (OD_600_ = 0.0001) after 10 hours of 4°C treatme*n*t (down), 22°C as control (up). Photos were taken after 48 h pathogen inoculation. (B) Bacterial populations in leaves described in (A). Values are displayed as mean ± s.e.m (*n* = 4 biological replicates). (C) ChIP-qPCR analysis of the *tga3^CR^* mutation impaired the relative enrichment of ICE1 to the *PR1* promoter. Col-0, *ICE1*-overexpressing (*ICE1-Myc*) and *tga3^CR^ ICE1-Myc* plants were treated for 2 h of 4°C, 22°C as control. (D) ChIP-qPCR analyses of the relative enrichment of TGA3 to the *PR1* promoter enhanced by ICE1. Col-0, *TGA3*-overexpressing plants (*TGA3-HA*) and *TGA3-HA ICE1-Myc* (hybridized F1 generation) were treated for 2 h of 4°C, 22°C as control. Treated leaves were pooled for ChIP assays using anti-HA antibody, the *ACTIN2* untranslated region sequence (*pACTIN2*) as a negative control in (C) and (D). (E) Transient transcriptional activity assays showing that TGA3 enhances the transcriptional activation of ICE1 to *PR1*. (F) Transient transcriptional activity assays showing that activation of the *PR1* promoter by ICE1 is compromised in the *tga3^CR^* mutant. (G) Transient transcriptional activity assays showing that activation of the *PR1* promoter by ICE1 is diminished in the *ice1-2* mutant. Each biological replicate was from different leaves of more than 60 plants in (E-G). (H) Transient transcriptional activity assays showing that ICE1 enhances the transcriptional activation of NPR1-TGA3 complex to *PR1* expression. Values are displayed as mean ± s.e.m (*n* = 3 biological replicates) for (C-H). Different letters indicate statistically significant differences (two-way ANOVA, *P* < 0.05). Experiments were repeated at least three times with similar trends.

ICE1 acts as an essential node integrating SA-regulated and low temperature-enhanced immunity by direct enrichment on the promoter of *PR1* (Fig. 5). To further delineate the regulatory relationship between ICE1 and TGA3, we conducted ChIP assays to investigate if TGA3 promotes the enrichment of ICE1 on the promoter of *PR1 in vivo*. For this purpose, *tga3^CR^ ICE1-Myc* and *TGA3-HA ICE1-Myc* seedlings were treated at 4°C for 2 h. Seedlings grown at 22°C served as the control. As shown in Fig. 7C, the enrichment of ICE1 on the *PR1* promoter was decreased in the background of *tga3^CR^* compared with Col-0. The accumulation of TGA3 on the *PR1* promoter was increased in the background of *ICE1-Myc-OE* plants compared with Col-0, and was further enhanced under low temperature (Fig. 7D). Dual-luciferase reporter assays using Arabidopsis mesophyll protoplasts were conducted to validate the effect of TGA3 on the transcriptional activation activity of ICE1. The effector consisted of GFP, TGA3, and ICE1 under the control of *Pro35S*, and the reporter consisted of the *PR1* promoter fused to the *LUC* gene (Supplemental Fig. S6). *LUC* expression driven by the *PR1* promoter was induced to higher level when TGA3 and ICE1 were co-expressed than when GFP and ICE1 were co-expressed (Fig. 7E). In contrast, ICE1 induced *LUC* expression was compromised in the *tga3^CR^* mutant compared with Col-0 (Fig. 7F). These results indicate that the ability of ICE1 to activate the transcription of *PR1* is promoted by TGA3. Noticeably, the ability of TGA3 to increase the transcription of *PR1* was diminished in the *ice1-2* mutant, suggesting that ICE1 also stimulates the transcriptional activation activities of TGA3 (Fig. 7G). These results support the notion that ICE1 and TGA3 act cooperatively to mediate cold-enhanced plant immunity through increased enrichment on the *PR1* promoter to activate its expression in association with SA.

The NPR1-TGA3 complex has been well documented that plays an indispensable role in *PR1* expression. Given ICE1 interacts with both NPR1 and TGA3, we questioned whether ICE1 could enhance the transcriptional activities of NPR1-TGA3 module on *PR1* expression. To clarify the potential role of ICE1 during the complex, we conducted the dual-luciferase reporter assay. Noticeably, compared with NPR1 and TGA3 were co-expressed, *LUC* expression driven by the *PR1* promoter was induced higher when NPR1, TGA3 and ICE1 were co-expressed (Fig. 7H), implying that ICE1 promotes the function of NPR1-TGA3 in *PR1* expression regulating.

### ICE1 Does not Affect the Biosynthesis of SA

In Arabidopsis, isochorismate, the precursor of SA, is mainly converted from chorismate by *ICS1/SID2* that is induced by pathogen infection (Wildermuth et al., 2001; Peng et al., 2021). AvrPphB Susceptible 3 (PBS3), a member of the GH3 acyl-adenylate/thioester-forming enzyme family, catalyzes the conjugation of isochorismate to glutamate to produce the key SA biosynthetic intermediate, isochorismate-9-glutamate, that can decay into SA spontaneously (Rekhter et al., 2019; Torrens-Spence et al., 2019). To determine whether ICE1-mediated immunity through regulating SA biosynthesis, we tested expression levels of *ICS1/SID2* and *PBS3* in Col-0 and *ice1-2* mutant plants at both normal and cold temperature. As shown in Supplemental Fig. S10, the expression levels of *ICS1/SID2* and *PBS3* displays no obvious differences between Col-0 and *ice1-2* mutant plants at both normal and cold temperature, suggesting that ICE1 did not involving in SA biosynthesis regulating.

## DISCUSSION

In this study, we demonstrate a previously uncharacterized function of ICE1 as an essential node bridging cold stimulus and plant immunity. We found that ICE1 is involved in cold-activated immunity as well as SA-regulated immunity, as demonstrated by the fact that the *ice1-2* mutant exhibited enhanced susceptibility to pathogen infection, including severe pathogen-related water soaking symptoms and high pathogen proliferation, and decreased expression of *PR1* (Fig. 1 and Fig. 3). Most importantly, the coordinated function of cold- and SA-enhanced immunity was diminished in *ice1-2* mutant plants, suggesting that ICE1 is the crucial regulator linking these two cues (Fig. 5). The major underlying mechanism is that NPR1 associates with ICE1 to promote its ability to activate *PR1* transcription (Fig. 2 and Fig. 4). Additionally, TGA3 works synergistically with ICE1 under low temperature (Fig. 6 and Fig. 7). The relationship between cold stress and immunity has not been fully elucidated in plants (Wigge, 2013; Kim et al., 2017; Wu et al., 2019). The identification of the NPR1-TGA3-ICE1 regulatory module in this study represents a significant step in understanding SA signaling during cold-activated resistance of plants to pathogen attack.

NPR1, the master regulator of SA signaling, is required for basal and systemically acquired resistance in plants (Cao et al., 1997; Kesarwani et al., 2007). Because it lacks a typical DNA-binding domain, NPR1 was long believed to function as a transcriptional coactivator that interacts with other proteins, including TGAs, EIN3, and Heat shock transcription factor 1 (HSFA1), to modulate physiological processes in plants (Zhang et al., 1999; Després et al., 2000; Olate et al., 2018; Huang et al., 2020; Wang et al., 2021; Yu et al., 2021). In terms of plant immune regulation, WRKYs and TGAs are well known to be direct targets of NPR1 and to regulate *PR1* gene expression (Zhang et al., 1999; Després et al., 2000; Zhou et al., 2000; Johnson et al., 2003; Wang et al., 2006; Saleh et al., 2015). Interestingly, in the present study, we demonstrate a novel function of ICE1 to activate SA signaling against pathogen infection. ICE1 interacts with NPR1 (Fig. 2) and functions in the same signal pathway with it (Fig. 4). As a MYC-like bHLH transcription factor, ICE1 can bind directly to the MYC recognition sequences of the *PR1* gene promoter to activate its expression (Supplemental Fig. S5; Fig. 3G-I), and its transcriptional activation activity is enhanced by NPR1 and TGA3 (Fig. 4C–F and Fig. 7C–F). In addition, ICE1 promotes the transcriptional activation of NPR1-TGA3 complex on expression of *PR1* (Fig. 7H). These findings not only provide evidence that ICE1 is an essential component of SA signal pathway at low temperature (Supplemental Fig. S11), but also increase our understanding of the role of the NPR1-ICE1-TGA3 regulatory module in integrating low temperature signal in activating plant immunity.

Recently, increasing attention has been paid to the interactions between low temperature stress and plant immunity. Previous studies have shown that the SA content increases in Arabidopsis plants exposed to cold stress (4°C) for more than 1 week, and that prolonged exposure (3 weeks) to low temperature promotes immunity against the bacterial plant pathogen, *Pst* DC3000 (Kim et al., 2013). Further study has clarified that low temperature can overcome the calmodulin-binding transcriptional activator 3 (CAMTA3) mediated genes inhibition of the SA pathway (Kim et al., 2017). Similarly, moderate low temperature (16°C) also enhances immunity accompanied by up-regulation of SA biosynthesis and signaling genes (Li et al., 2020). Wu et al. found that exposure to short-term cold stress (4°C for 10 hours) activates SA-dependent immunity as well as H_2_O_2_ and callose deposition (Wu et al., 2019). The results of those studies imply that SA may play a crucial role in low temperature-activated immunity, however, the underlying mechanism remained poorly characterized. In this study, we discovered that ICE1 acts an indispensable regulatory node bridging low temperature cues and the SA signaling pathway. Our results show that low temperature treatment enhances disease resistance against the bacterial pathogen, *Pst* DC3000 (Fig. 1A–C), and that ICE1 is required for this modulation (Fig. 1E–H). Interestingly, we found that cold stress coordinates SA signaling to promote immunity, and this synergistic regulation occurs in an ICE1-dependent manner. This is highlighted by the fact that the enhanced immunity caused by low temperature combined with SA signaling was compromised in *ice1-2* mutant plants compared with Col-0 plants (Fig. 5A, B). Low temperature has multifaceted influences on the SA signaling pathway, which involves ICE1. For instance, in the Co-IP assays, low temperature facilitated interactions between ICE1 and NPR1 (Fig. 2D, E). Notably, in the BiFC and LUC assays, the NPR1 and ICE1 interaction only occurred after cold treatment (Fig. 2B, C), consistent with the results of previous studies showing that NPR1 accumulates in response to low temperature and enters the nucleus as a monomer to exert its function as a co-factor (Olate et al., 2018). The enrichment of ICE1–TGA3 on the *PR1* promoter and the ability of this complex to transcriptionally activate *PR1* were found to be significantly enhanced under low temperature (Fig. 7C–G). These results provide direct evidence for the co-operative regulation of low temperature cues and SA signaling *via* ICE1, illuminating the relationship between cold stress and SA-regulated immunity.

It has been proposed that NPR1 is present in both an inactive oligomer form and active monomer form and that SA treatment and low temperature promote the conformation change from oligomer to monomer (Mou et al., 2003; Tada et al., 2008; Yu et al., 2022). Although, in normal temperature and absence of SA, the majority of NPR1 proteins are in the oligomer form, some NPR1 proteins are in the monomer form, which may explain why NPR1 can interact with other proteins, including HSFA1, GID1 and EIN3 in normal temperature and absence of SA, however, the interactions can be enhanced by SA and low temperature (Olate et al., 2018; Huang et al., 2020; Yu et al., 2022). Consistent with these findings, we found that NPR1 could interact with ICE1 in normal temperature and facilitate its transcriptional activation on *PR1* expression (Fig. 4). Interestingly, association of NPR1 and ICE1 was enhanced by low temperature (Fig. 2B-E). Low temperature may promote the accumulation of NPR1 monomer in the nuclei, leading to more active NPR1 proteins targeting ICE1.

Temperature is an essential environmental factor affecting plant growth and development, as well as resistance against pathogens. Recent studies have shown that elevated temperatures negatively affect pathogen resistance by suppressing SA production in Arabidopsis (Huot et al., 2017; Kim et al., 2022). Interestingly, the results of several studies, including ours, show that the SA pathway also plays indispensable role in the defense against bacterial pathogen attack under cold stress conditions (Fig. 5) (Seo et al., 2010; Kim et al., 2013; Kim et al., 2017; Wu et al., 2019; Li et al., 2020). Climate change will have a profound effect on plant immunity, so an in-depth understanding of the regulatory role of low temperature on plant immunity is instrumental for revealing the mechanisms of environmental adaptation in plants under cold stress. The results of studies on the temperature regulation of pathogen resistance may provide a theoretical basis for the development of new strategies to improve plants’ resistance to pathogens in future agricultural production. In addition, because of their sessile lifestyle, plants are constantly exposed to a variety of abiotic stresses, and endogenous hormone signals often interact with multiple exogenous environmental cues to regulate their resistance to pathogens. Dissecting the regulatory mechanism of the integration of low temperature and SA signals in response to pathogen attack also provides new possibilities for research on the cross-talk between SA signaling and other environmental signals.

## MATERIALS AND METHODS

### Plant Materials and Growth Conditions

The wild-type and mutant Arabidopsis plants used in this study were in the Columbia (Col-0) genetic background. All Arabidopsis plants were soil-grown for four weeks under an artificial growth chamber at 22°C and 60% relative humidity, and 10 h light/14 h dark photoperiod. The *ice1-2* (SALK_003155; (Kanaoka et al., 2008)) mutant and *npr1-1* (CS3726; (Cao et al., 1994)) mutant have been described. The same *cbfs* triple mutants with study of Jia et al. (2016) was used in this study (Jia et al., 2016). Seeds of *ice1-2/ICE1* were kindly provided by Prof. Shuhua Yang (China Agricultural University). The *35S::GFP-ICE1* (CS68099) line was obtained from the Arabidopsis Resource Center at Ohio State University (http://abrc.osu.edu). To generate the overexpression transgenic plants, the full-length cDNA of *ICE1, TGA3* were cloned into the pOCA30 vector by *BamHI* (Thermo Scientific) and *SalI* (Thermo Scientific) in the sense orientation behind the CaMV 35S promoter using T4 DNA ligase (Thermofisher), respectively. Primers used for transgenic construction are listed in Supplemental Data Set 1. For CRISPR/Cas9-mediated editing of *TGA3*, one guide RNA was designed by CRISPR-P 2.0 (http://crispr.hzau.edu.cn/CRISPR2/) to target the third exon of *TGA3*. This guide RNA, driven by the AtU6a promoter, was cloned into the pMH-SA binary vector carrying Cas9 by *SpeI* (Thermo Scientific) and *AscI* (Thermo Scientific) using T4 DNA ligase (Thermofisher) (Liang et al., 2016). The detailed information about guide sequence and direct sequencing of PCR products containing targeted sites of *TGA3* in Arabidopsis plants are showed in the Supplemental Fig. S9. The genotyping/sequencing primers are listed in Supplemental Data Set 1.

For low temperature treatment, plants were grown for 4 weeks at 22°C, followed by 3 weeks or 10 h of 4°C treatment. Plants had roughly 20 rosette leaves and not yet reached to the flowering stage before cold treatment. Control plants were directly inoculated with *Pst* DC3000, and cold-treated plants were transferred to a cold incubator (4°C, 65% relative humidity, 10 h light/14 h dark photoperiod) for 3 weeks or 10 h treatment before pathogen inoculation.

For the combined cold and BTH treatment, 4-week-old plants sprayed with mock and BTH and kept in a growth chamber with normal temperature (22°C) for 14 h. After then, the low temperature experimental plants were transferred to a low temperature (4°C) growth chamber for 10 h cold treatment, while control plants still kept in the growth chamber with normal temperature (22°C) for additional 10 h. In this way, the plants can suffer from different treatment conditions, such as 24 h mock and 24 h BTH with normal temperature as well as 24 h mock and 24 h BTH with 10 h low temperature before *Pst* DC3000 inoculation.

### Bacterial Disease Assays

Syringe-infiltration was performed in this study. Briefly, *Pst* DC3000 was cultured in modified Luria–Bertani medium (Xin et al., 2016) containing 100 mg l^−1^ rifampicin at 28 °C to an OD_600_ of 0.8–1.0. Bacteria were collected by centrifugation and re-suspended in 0.25 mM MgCl_2_. Cell density was adjusted to OD_600_ = 0.2 (approximately 1 × 10^8^ cfu ml^−1^) and further diluted to cell densities of 1 × 10^5^–1 × 10^6^ cfu ml^−1^. The infiltrated plants were first placed under ambient humidity for 1–2 h to allow water to evaporate, and then placed in high humidity (about 95%; by covering plants with a clear plastic dome) for disease development after the leaves returned to the state before infiltration.

For quantification of *Pst* DC3000 bacterial populations, leaf disks were taken using a disposable biopsy punch with 4 mm diameter (Integra LifeSciences) and ground in 0.25 mM MgCl_2_. Colony-forming units were determined by serial dilutions and plating on Luria Marine plates containing 100 mg l^−1^ rifampicin. One biological replicate consists of 12 leaf disks (that is, from 3 leaves of one plant) and at least 3 biological replicates were included in each experiment. Experiments were repeated at least three times.

For BTH protection assay, four-week-old *Arabidopsis thaliana* were sprayed with benzo_(1,2,3) thiadiazole-7-carbothioic acid-S-methyl ester (BTH; Chem Service, 0.1% DMSO, 0.05% Silwet L77) or Mock (0.1% DMSO, 0.05% Silwet L77) 24 h before bacterial syringe-infiltration.

### RNA Extraction and RT-qPCR

Total RNA was extracted by using Trizol reagent (Invitrogen). The RT-qPCR analysis was conducted as described (He et al., 2023; Li et al., 2023). In brief, 1.0 μg DNase-treated RNA was reverse transcribed in a reaction volume of 20 μl containing oligo-(dT)19 primer and Moloney murine leukemia virus reverse transcriptase (Thermo Fisher Scientific). RT-qPCR analysis was then performed with 1.0 μl of 5-fold diluted cDNA using a SYBR premixed Ex Taq kit (Takara), and a Light Cycler 480 Real-Time PCR System (Roche) for the reaction. In each RT-qPCR experiment, at least three biological replicates were used and three technical replicates were performed for each biological replicate. The *ACTIN2* (*AT3G18780*) gene was used as an internal reference gene.

### Electrophoresis Mobility Shift Assay (EMSA)

To generate the ICE1-His recombinant protein, the sequence encoding full-length *ICE1* was cloned into pET28a using ClonExpress II One Step Cloning Kit (Vazyme) and transformed into *Escherichia coli* strain BL21 (DE3) (TIANGEN). The protein was induced with 0.5 mM isopropyl β-D-1-thiogalactopyranoside (IPTG) for 18 h at 16°C, and purified with Ni-NTA resin. Oligonucleotide probes containing wild type E-box were synthesized and labeled with 5 ’ Biotin modification (BGI, China). The unlabeled wild type E-box served as a competitor, and the mutated E-box modified by biotin served as a negative control. Double-stranded probes were generated by annealing of forward/reverse primers. EMSA was performed using biotin-labeled probes and Chemiluminescent EMSA Kit (Beyotime; GS009) referring to the instructions. For the binding reaction, ICE1-His protein was incubated with binding buffer containing 1 μl of biotin-labeled oligonucleotide in a total volume of 10 μl. Competition experiments were executed using 50-fold and 200-fold unlabeled double-stranded DNA. The DNA-protein complexes were separated on a 6% polyacrylamide gel in 0.5 × Tris–borate-EDTA buffer. The primers used for vector construction are listed in Supplemental Data Set 1.

### Yeast Two-Hybrid Screening and Confirmation

The full-length coding sequences (CDS) of *NPR1* and *TGA* were fused to the bait vector pGBKT7 (Clontech, Palo Alto, CA, USA) by *NdeI* (Thermo Scientific) and *EcoRI* (Thermo Scientific) using ClonExpress II One Step Cloning Kit (Vazyme) to generate BD-NPR1 and BD-TGA, while the full-length coding sequence of *ICE1* was ligated to the prey vector pGADT7 to produce AD-ICE1. To identify structural domains essential for protein interactions, several truncated NPR1 and TGA3 were cloned into pGBKT7 and truncated ICE1 into pGADT7. The yeast two hybrid (Y2H) assays were performed as described previously (Li et al., 2023). The vector pairs were co-transformed into the yeast strain AH109, protein interactions were indicated by the ability of cells to grow on dropout medium lacking Leu, Trp, His, and Ade for 2 days after plating. The primers used for vector construction are listed in Supplemental Data Set 1.

### Split Luciferase Complementation Assay

The split luciferase complementation assays were performed as described previously (Fujikawa and Kato, 2007; Chen et al., 2008). The full-length CDS of *ICE1* was cloned into the pCAMBIA1300-nLUC vector by *BamHI* (Thermo Scientific) and *SalI* (Thermo Scientific) under the control of the 35S promoter using ClonExpress II One Step Cloning Kit (Vazyme). Full-length CDS of NPR1 and TGA3 were cloned into the pCAMBIA1300-cLUC vectors by *KpnI* (Thermo Scientific) and *SalI* (Thermo Scientific) using ClonExpress II One Step Cloning Kit (Vazyme), respectively. After co-expression in *Nicotiana benthamiana* leaves for 48 h, the leaves were sprayed with 1 mM D-luciferin (0.01% Triton X-100) solution and then incubated for 5 min in the dark before detecting. Luciferase intensity was visualized on Tanon-5200 Chemiluminescent Imaging System (Tanon Science&Technology). The primers used for vector construction are listed in Supplemental Data Set 1.

### Bimolecular Fluorescence Complementation Assays

The sequences encoding the C-terminal (64 amino acids) YFP (cYFP) fragment and the N-terminal 173 amino acid-enhanced YFP (nYFP) fragment were amplified by PCR and cloned into the labeled pFGC5941 plasmid to generate pFGC-nYFP and pFGC-cYFP, respectively (Kim et al., 2008). The full-length or truncated CDS sequences of *ICE1* were cloned into pFGC-nYFP by *BamHI* (Thermo Scientific) using ClonExpress II One Step Cloning Kit (Vazyme) to generate the N-terminal in-frame fusions with n-YFP (ICE1-nYFP, ICE1^1-260^-nYFP and ICE1^261-494^-nYFP), whereas full-length or truncated *NPR1* and *TGA* coding sequences were inserted into pFGC-cYFP by *BamHI* (Thermo Scientific) using ClonExpress II One Step Cloning Kit (Vazyme) to form a C-terminal intraframe fusion (NPR1-cYFP, NPR1^1-194^-cYFP, NPR1^178-593^-cYFP and TGA3-cYFP). The plasmids were transformed into *Agrobacterium tumefaciens* strain GV3101 and injected into tobacco (*N. benthamiana*) leaves. As described previously (He et al., 2023; Li et al., 2023), YFP and 4,6-diamidino-2-phenylindole (DAPI) fluorescence of infected leaves were observed under a confocal laser scanning microscope (Olympus) after 48 h of co-expression. Primers used for cloning are listed in Supplemental Data Set 1.

### Co-immunoprecipitation Assays

To generate proteins fused with YFP or Myc tags, we amplified the full-length CDS of *NPR1* and *TGA3* and recombined into the intermediate vector pDonor207 using the BP recombination kit (Invitrogen), then cloned into the destination vector pEarleyGate101 with YFP-tagged sequences under the control of *Pro35S* using the LR recombination kit (Invitrogen) to obtain 35S::NPR1-YFP and 35S::TGA3-YFP constructs, whereas the full-length CDS of *ICE1* was inserted into the destination vector Myc-tagged pGWB517 using the same method to get 35S::ICE1-Myc. Arabidopsis protoplasts were transformed with different combinations of plasmids, including 35S::YFP and 35S:: ICE1-Myc, 35S::NPR1-YFP and 35S::ICE1-Myc, 35S::TGA3-YFP and 35S:ICE1-Myc at 22°C for 16 h according to the Sheen laboratory protocol (Sheen, 2001). Total protein was extracted with an IP buffer containing 50 mM MOPs, 5 mM EDTA, 0.2% v/v Triton X-100, 5 mM dithiothreitol, 1 mM PMSF and 1 x complete protease inhibitor cocktail (Roche). Immunoprecipitation experiments with GFP-trap beads were performed according to the manufacturer’s protocol. Briefly, cell lysates were incubated with GFP-trap beads (ChromoTek) at 4°C overnight. After incubation, the beads were washed three to five times with IP buffer, and then co-immunoprecipitated proteins were detected by immunoblotting with an anti-Myc antibody (Abmart; 1:10,000). Primers used for vector construction are listed in Supplementary Data Set 1.

For Co-IP assay using stable transgenic seedlings, total proteins were extracted with IP buffer from *35S:HA-NPR1*, *35S:ICE1-Myc* and *35S:HA-NPR1 35S:ICE1-Myc* transgenic Arabidopsis seedlings at both normal temperature and 2 h cold treatments, Col-0 seedlings used as negative control. Proteins immunoprecipitated with Myc-trap beads (ChromoTek) at 4°C overnight, then the beads were washed three to five times with IP buffer. The Co-immunoprecipitated proteins were detected with anti-HA antibody (ZENBIO; 1:10,000).

### Chromatin Immunoprecipitation

Chromatin immunoprecipitation (ChIP) Assays were performed substantially as described earlier (Mukhopadhyay et al., 2008; Jiang et al., 2014). In brief, Col-0, *ICE1-Myc*, *npr1-1 ICE1-Myc*, *NPR1-HA ICE1-Myc*, *TGA3-HA ICE1-Myc*, and *tga3^CR^ ICE1-Myc* plants were treated with 1% formaldehyde (cross-linking treatment) to isolate their chromatin. Proteins were immunoprecipitated using anti-Myc antibody (1:1,000). The precipitated DNA was purified using a PCR purification kit (Qiagen). To quantitatively determine the ICE1–DNA (target promoter) binding, qPCR analysis was performed with the *ACTIN2* 3’ untranslated region sequence as an endogenous control. Relative enrichment was calculated based on DNA binding ratio. Primers used for the ChIP-qPCR are listed in Supplemental Data Set 1.

### Transient Transcriptional Activation Assays

The full-length CDS of *NPR1* was cloned into the pGreenII 62-SK vector by *SacI* (Thermo Scientific) and *BamHI* (Thermo Scientific), while full-length CDS of *ICE1*, *TGA3*, and *GFP* were cloned into the pGreenII 62-SK vector by *BamHI* (Thermo Scientific) and *EcoRI* (Thermo Scientific) using T4 DNA ligase (Thermofisher) as effectors, respectively. The promoter sequence of *PR1* was amplified by PCR and inserted into the pGreenII 0800-LUC vector (Biovector Science Lab) by *SacI* (Thermo Scientific) and *BamHI* (Thermo Scientific) using T4 DNA ligase (Thermofisher) as the reporter (Hellens et al., 2005). Combinations of plasmids were transformed into the Col-0 and/or mutant leaf mesophyll protoplasts according to the Sheen laboratory protocol (Sheen, 2001). Transfected cells were cultured for 16-18 h followed by analysis of relative luciferase (LUC) activity using the Dual-Luciferase Reporter Assay System (Promega), which detects the activities of firefly LUC and the internal control Renilla reniformis (REN). Primers used for vector construction are listed in Supplemental Data Set 1.

## STATISTICAL ANALYSIS

Statistical analysis was performed by analysis of variance. The results of statistical analyses are shown in Supplemental Data Set S2.

## ACCESSION NUMBERS

*Arabidopsis* Genome Initiative numbers for the genes discussed in this article are as follows: *ICE1*, AT3G26744; *NPR1*, AT1G64280; *TGA1*, AT5G65210; *TGA2*, AT5G06950; *TGA3*, AT1G22070; *TGA4*, AT5G10030; *TGA*5, AT5G06960; *TGA6*, AT3G12250; *TGA7*, AT1G77920; *PR1*, AT2G14610; *PR2*, AT3G57260; *PR5*, AT1G75040; *WRKY70*, AT3G56400; *WRKY18*, AT4G31800; *WRKY38*, AT5G22570; *WRKY62*, AT5G01900; *ICS1*(SID2), AT1G74710; *PBS3*, AT5G13320; *ACTIN2*, AT3G18780.

## Supporting information

Supplemental Information-Version 2

## ACKNOWLEDGMENTS

We thank Sheng Yang He for his thoughtful revision and constructive comments on the manuscript. We would like to thank our lab members Diqiu Yu, Houping Wang and Pengcheng Wang for their suggestions during this work. We also thank Jennifer Smith, PhD, from Liwen Bianji (Edanz) (www.liwenbianji.cn/) for editing the English text of a draft of this manuscript.

## AUTHOR CONTRIBUTIONS

S.Q.L. and Y.J.J. designed and performed experiments; S.Q.L., L.H., Y.P.Y. Y.X.Z. and X.H. and Y.R.H analyzed data; S.Q.L. and Y.J.J. wrote the manuscript. All authors read and approved the final manuscript.

## SUPPLEMENTAL DATA

The following materials are available in the online version of this article.

**Supplemental Figure S1.** No interacting signal of ICE1 and NPR1 was detected at normal temperature.

**Supplemental Figure S2.** Percentage of transformed cells with YFP fluorescence in the BiFC assays presented in Figures 2B and 6B.

**Supplemental Figure S3.** Transcriptional and protein levels of ICE1 responding to BTH treatment.

**Supplemental Figure S4.** Expression of SA-related defense genes.

**Supplemental Figure S5.** The promoter sequence of *PR1*.

**Supplemental Figure S6.** Schematic of effectors and reporters used in transient transactivation assays.

**Supplemental Figure S7.** *CBF* genes does not mediate low temperature-enhanced and SA-regulated immunity.

**Supplemental Figure S8.** The bHLH motif and bZIP motif was required for interaction of ICE1 and TGA3.

**Supplemental Figure S9.** Generation of *tga3^CR^* knockout mutant by CRISPR/Cas9 technology.

**Supplemental Figure S10.** Expression of *ICS1/SID2* and *PBS3* in Col-0 and *ice1-2* mutant plants.

**Supplemental Figure S11**. Working model of NPR1-TGA3/ICE1 regulating module against *Pst* DC3000 after cold stress.

**Supplemental Figure S12.** Uncropped images for protein gels.

**Supplemental Data Set 1.** Primers used in the study.

**Supplemental Data Set 2.** The results of statistical analyses performed by ANOVA.

## FUNDING

This study was supported by funding from the Natural Science Foundation of China (Grants 32360082 to Y.J.J.) and the Young and Middle Aged Academic and technical Leaders of Yunnan Province (202105AC160028 to Y.J.J.).

*Conflict of interest statement.* All authors state that they have no conflict of interest in relation to this research.

## DATA AVAILABILITY

All data supporting the findings of this study are available within the article and its supplemental materials.

## Notes

### Competing Interest Statement

The authors have declared no competing interest.

### Summary of Updates

Main figures and supplemental figures are revised.

## REFERENCES

1. An, J.P., Wang, X.F., Zhang, X.W., You, C.X., and Hao, Y.J. (2021). Apple B-box protein BBX37 regulates jasmonic acid mediated cold tolerance through the JAZ-BBX37-ICE1-CBF pathway and undergoes MIEL1-mediated ubiquitination and degradation. New Phytol 229, 2707–2729.

2. An, J.P., Xu, R.R., Liu, X., Su, L., Yang, K., Wang, X.F., Wang, G.L., and You, C.X. (2022). Abscisic acid insensitive 4 interacts with ICE1 and JAZ proteins to regulate ABA signaling-mediated cold tolerance in apple. J Exp Bot 73, 980–997.

3. Benedict, C., Geisler, M., Trygg, J., Huner, N., and Hurry, V. (2006). Consensus by democracy. Using meta-analyses of microarray and genomic data to model the cold acclimation signaling pathway in Arabidopsis. Plant Physiol 141, 1219–1232.

4. Bernsdorff, F., Döring, A.C., Gruner, K., Schuck, S., Bräutigam, A., and Zeier, J. (2016). Pipecolic acid orchestrates plant systemic acquired resistance and defense priming via salicylic acid-dependent and - independent pathways. Plant Cell 28, 102–129.

5. Bi, G., Su, M., Li, N., Liang, Y., Dang, S., Xu, J., Hu, M., Wang, J., Zou, M., Deng, Y., Li, Q., Huang, S., Li, J., Chai, J., He, K., Chen, Y.H., and Zhou, J.M. (2021). The ZAR1 resistosome is a calcium-permeable channel triggering plant immune signaling. Cell 184, 3528–3541.

6. Boller, T., and Felix, G. (2009). A renaissance of elicitors: perception of microbe-associated molecular patterns and danger signals by pattern-recognition receptors. Annu Rev of Plant Biol 60, 379–406.

7. Cao, H., Bowling, S.A., Gordon, A.S., and Dong, X. (1994). Characterization of an Arabidopsis mutant that is nonresponsive to inducers of systemic acquired resistance. Plant Cell 6, 1583–1592.

8. Cao, H., Glazebrook, J., Clarke, J.D., Volko, S., and Dong, X. (1997). The Arabidopsis *NPR1* gene that controls systemic acquired resistance encodes a novel protein containing ankyrin repeats. Cell 88, 57–63.

9. Chen, H., Zou, Y., Shang, Y., Lin, H., Wang, Y., Cai, R., Tang, X., and Zhou, J.M. (2008). Firefly luciferase complementation imaging assay for protein-protein interactions in plants. Plant Physiol 146, 368–376.

10. Chen, H., Raffaele, S., and Dong, S. (2021). Silent control: microbial plant pathogens evade host immunity without coding sequence changes. FEMS Microbiol Rev 45, 1–16.

11. Chinnusamy, V., Zhu, J., and Zhu, J.K. (2007). Cold stress regulation of gene expression in plants. Trends Plant Sci 12, 444–451.

12. Chinnusamy, V., Ohta, M., Kanrar, S., Lee, B.H., Hong, X., Agarwal, M., and Zhu, J.K. (2003). ICE1: a regulator of cold-induced transcriptome and freezing tolerance in Arabidopsis. Genes Dev 17, 1043–1054.

13. Choi, J., Huh, S.U., Kojima, M., Sakakibara, H., Paek, K.H., and Hwang, I. (2010). The cytokinin-activated transcription factor ARR2 promotes plant immunity via TGA3/NPR1-dependent salicylic acid signaling in Arabidopsis. Dev Cell 19, 284–295.

14. Couto, D., and Zipfel, C. (2016). Regulation of pattern recognition receptor signalling in plants. Nat Rev Immunol 16, 537–552.

15. Deslandes, L., and Rivas, S. (2012). Catch me if you can: bacterial effectors and plant targets. Trends Plant Sci 17, 644–655.

16. Després, C., DeLong, C., Glaze, S., Liu, E., and Fobert, P.R. (2000). The Arabidopsis NPR1/NIM1 protein enhances the DNA binding activity of a subgroup of the TGA family of bZIP transcription factors. Plant Cell 12, 279–290.

17. Ding, Y., Li, H., Zhang, X., Xie, Q., Gong, Z., and Yang, S. (2015). OST1 kinase modulates freezing tolerance by enhancing ICE1 stability in Arabidopsis. Dev Cell 32, 278–289.

18. Ding, Y., Sun, T., Ao, K., Peng, Y., Zhang, Y., Li, X., and Zhang, Y. (2018). Opposite roles of salicylic acid receptors NPR1 and NPR3/NPR4 in transcriptional regulation of plant immunity. Cell 173, 1454–1467.e1415.

19. Dodds, P.N., and Rathjen, J.P. (2010). Plant immunity: towards an integrated view of plant-pathogen interactions. Nature Rev Genet 11, 539–548.

20. Dong, C.H., Agarwal, M., Zhang, Y., Xie, Q., and Zhu, J.K. (2006). The negative regulator of plant cold responses, HOS1, is a RING E3 ligase that mediates the ubiquitination and degradation of ICE1. Proc Natl Acad Sci USA 103, 8281–8286.

21. Durrant, W.E., and Dong, X. (2004). Systemic acquired resistance. Annu Rev Phytopathol 42, 185–209.

22. Fujikawa, Y., and Kato, N. (2007). Split luciferase complementation assay to study protein-protein interactions in Arabidopsis protoplasts. Plant J 52, 185–195.

23. Gilmour, S.J., Zarka, D.G., Stockinger, E.J., Salazar, M.P., Houghton, J.M., and Thomashow, M.F. (1998). Low temperature regulation of the Arabidopsis CBF family of AP2 transcriptional activators as an early step in cold-induced *COR* gene expression. Plant J 16, 433–442.

24. Griffith, M., and Yaish, M.W. (2004). Antifreeze proteins in overwintering plants: a tale of two activities. Trends Plant Sci 9, 399–405.

25. Han, Q., Tan, W., Zhao, Y., Yang, F., Yao, X., Lin, H., and Zhang, D. (2022). Salicylic acid-activated BIN2 phosphorylation of TGA3 promotes Arabidopsis *PR* gene expression and disease resistance. EMBO J, e110682.

26. Hartmann, M., and Zeier, J. (2019). N-hydroxypipecolic acid and salicylic acid: a metabolic duo for systemic acquired resistance. Curr Opin Plant Biol 50, 44–57.

27. He, K., Du, J., Han, X., Li, H., Kui, M., Zhang, J., Huang, Z., Fu, Q., Jiang, Y., and Hu, Y. (2023). PHOSPHATE STARVATION RESPONSE1 (PHR1) interacts with JASMONATE ZIM-DOMAIN (JAZ) and MYC2 to modulate phosphate deficiency-induced jasmonate signaling in Arabidopsis. Plant Cell 35, 2132–2156.

28. Hellens, R.P., Allan, A.C., Friel, E.N., Bolitho, K., Grafton, K., Templeton, M.D., Karunairetnam, S., Gleave, A.P., and Laing, W.A. (2005). Transient expression vectors for functional genomics, quantification of promoter activity and RNA silencing in plants. Plant Methods 1, 13.

29. Hu, Y., Han, X., Yang, M., Zhang, M., Pan, J., and Yu, D. (2019). The transcription factor INDUCER OF CBF EXPRESSION1 interacts with ABSCISIC ACID INSENSITIVE5 and DELLA proteins to fine-tune abscisic acid signaling during seed germination in Arabidopsis. Plant Cell 31, 1520–1538.

30. Huang, P., Dong, Z., Guo, P., Zhang, X., Qiu, Y., Li, B., Wang, Y., and Guo, H. (2020). Salicylic acid suppresses apical hook formation via NPR1-mediated repression of EIN3 and EIL1 in Arabidopsis. Plant Cell 32, 612–629.

31. Huang, X., Li, J., Bao, F., Zhang, X., and Yang, S. (2010). A gain-of-function mutation in the Arabidopsis disease resistance gene *RPP4* confers sensitivity to low temperature. Plant Physiol 154, 796–809.

32. Huot, B., Castroverde, C.D.M., Velasquez, A.C., Hubbard, E., Pulman, J.A., Yao, J., Childs, K.L., Tsuda, K., Montgomery, B.L., and He, S.Y. (2017). Dual impact of elevated temperature on plant defence and bacterial virulence in Arabidopsis. Nat Commun 8, 1808.

33. Jia, X., Wang, L., Zhao, H., Zhang, Y., Chen, Z., Xu, L., and Yi, K. (2023). The origin and evolution of salicylic acid signaling and biosynthesis in plants. Mol Plant 16, 245–259.

34. Jia, Y., Ding, Y., Shi, Y., Zhang, X., Gong, Z., and Yang, S. (2016). The *cbfs* triple mutants reveal the essential functions of CBFs in cold acclimation and allow the definition of CBF regulons in Arabidopsis. New Phytol 212, 345–353.

35. Jiang, Y.J., Liang, G., Yang, S.Z., and Yu, D.Q. (2014). Arabidopsis WRKY57 functions as a node of convergence for jasmonic acid- and auxin-mediated signaling in jasmonic acid-induced leaf senescence. Plant Cell 26, 230–245.

36. Johnson, C., Boden, E., and Arias, J. (2003). Salicylic acid and NPR1 induce the recruitment of trans-activating TGA factors to a defense gene promoter in Arabidopsis. Plant Cell 15, 1846–1858.

37. Jones, J.D., and Dangl, J.L. (2006). The plant immune system. Nature 444, 323–329.

38. Kachroo, P., Liu, H., and Kachroo, A. (2020). Salicylic acid: transport and long-distance immune signaling. Curr Opin Virol 42, 53–57.

39. Kanaoka, M.M., Pillitteri, L.J., Fujii, H., Yoshida, Y., Bogenschutz, N.L., Takabayashi, J., Zhu, J.K., and Torii, K.U. (2008). SCREAM/ICE1 and SCREAM2 specify three cell-state transitional steps leading to Arabidopsis stomatal differentiation. Plant Cell 20, 1775–1785.

40. Kesarwani, M., Yoo, J., and Dong, X. (2007). Genetic interactions of TGA transcription factors in the regulation of pathogenesis-related genes and disease resistance in Arabidopsis. Plant Physiol 144, 336–346.

41. Kim, J.H., Castroverde, C.D.M., Huang, S., Li, C., Hilleary, R., Seroka, A., Sohrabi, R., Medina-Yerena, D., Huot, B., Wang, J., Nomura, K., et al. (2022). Increasing the resilience of plant immunity to a warming climate. Nature 607, 339–344.

42. Kim, K.C., Lai, Z., Fan, B., and Chen, Z. (2008). Arabidopsis WRKY38 and WRKY62 transcription factors interact with histone deacetylase 19 in basal defense. Plant Cell 20, 2357–2371.

43. Kim, Y., Park, S., Gilmour, S.J., and Thomashow, M.F. (2013). Roles of CAMTA transcription factors and salicylic acid in configuring the low-temperature transcriptome and freezing tolerance of Arabidopsis. Plant J 75, 364–376.

44. Kim, Y.S., Lee, M., Lee, J.H., Lee, H.J., and Park, C.M. (2015). The unified ICE-CBF pathway provides a transcriptional feedback control of freezing tolerance during cold acclimation in Arabidopsis. Plant Mol Biol 89, 187–201.

45. Kim, Y.S., An, C., Park, S., Gilmour, S.J., Wang, L., Renna, L., Brandizzi, F., Grumet, R., and Thomashow, M.F. (2017). CAMTA-mediated regulation of salicylic acid immunity pathway genes in Arabidopsis exposed to low temperature and pathogen infection. Plant Cell 29, 2465–2477.

46. Nomura, K., He, S.Y. (2005). Powerful screens for bacterial virulence proteins. Proc Natl Acad Sci USA 102, 3527–3528.

47. Kumar, S., Zavaliev, R., Wu, Q., Zhou, Y., Cheng, J., Dillard, L., Powers, J., Withers, J., Zhao, J., Guan, Z., Borgnia, M.J., Bartesaghi, A., Dong, X., and Zhou, P. (2022). Structural basis of NPR1 in activating plant immunity. Nature 605, 561–566.

48. Lee, B.H., Henderson, D.A., and Zhu, J.K. (2005). The Arabidopsis cold-responsive transcriptome and its regulation by ICE1. Plant Cell 17, 3155–3175.

49. Lee, J.H., Jung, J.H., and Park, C.M. (2015). INDUCER OF CBF EXPRESSION 1 integrates cold signals into FLOWERING LOCUS C-mediated flowering pathways in Arabidopsis. Plant J 84, 29–40.

50. Lee, J.H., Jung, J.H., and Park, C.M. (2017). Light inhibits COP1-mediated degradation of ICE transcription factors to induce stomatal development in Arabidopsis. Plant Cell 29, 2817–2830.

51. Li, H., Ding, Y., Shi, Y., Zhang, X., Zhang, S., Gong, Z., and Yang, S. (2017). MPK3- and MPK6-mediated ICE1 phosphorylation negatively regulates ICE1 stability and freezing tolerance in Arabidopsis. Dev Cell 43, 630–642.e634.

52. Li, Z., Li, S., Jin, D., Yang, Y., Pu, Z., Han, X., Hu, Y., and Jiang, Y. (2023). U-box E3 ubiquitin ligase PUB8 attenuates abscisic acid responses during early seedling growth. Plant Physiol 191, 2519–2533.

53. Li, Z., Liu, H., Ding, Z., Yan, J., Yu, H., Pan, R., Hu, J., Guan, Y., and Hua, J. (2020). Low temperature enhances plant immunity via salicylic acid pathway genes that are repressed by ethylene. Plant Physiol 182, 626–639.

54. Liang, G., Zhang, H., Lou, D., and Yu, D. (2016). Selection of highly efficient sgRNAs for CRISPR/Cas9-based plant genome editing. Sci Rep 6, 21451.

55. Lopez, V.A., Park, B.C., Nowak, D., Sreelatha, A., Zembek, P., Fernandez, J., Servage, K.A., Gradowski, M., Hennig, J., Tomchick, D.R., Pawlowski, K., Krzymowska, M., and Tagliabracci, V.S. (2019). A bacterial effector mimics a host HSP90 client to undermine immunity. Cell 179, 205–218.

56. Ma, S., Lapin, D., Liu, L., Sun, Y., Song, W., Zhang, X., Logemann, E., Yu, D., Wang, J., Jirschitzka, J., Han, Z., Schulze-Lefert, P., Parker, J.E., and Chai, J. (2020). Direct pathogen-induced assembly of an NLR immune receptor complex to form a holoenzyme. Science 370, 1184–1193.

57. Macho, A.P., and Zipfel, C. (2015). Targeting of plant pattern recognition receptor-triggered immunity by bacterial type-III secretion system effectors. Curr Opin Microbiol 23, 14–22.

58. Miura, K., Jin, J.B., Lee, J., Yoo, C.Y., Stirm, V., Miura, T., Ashworth, E.N., Bressan, R.A., Yun, D.J., and Hasegawa, P.M. (2007). SIZ1-mediated sumoylation of ICE1 controls *CBF3/DREB1A* expression and freezing tolerance in Arabidopsis. Plant Cell 19, 1403–1414.

59. Mou, Z., Fan, W., and Dong, X. (2003). Inducers of plant systemic acquired resistance regulate NPR1 function through redox changes. Cell 113, 935–944.

60. Mukhopadhyay, A., Deplancke, B., Walhout, A.J., and Tissenbaum, H.A. (2008). Chromatin immunoprecipitation (ChIP) coupled to detection by quantitative real-time PCR to study transcription factor binding to DNA in Caenorhabditis elegans. Nat Protoc 3, 698–709.

61. Ngou, B.P.M., Ding, P., and Jones, J.D.G. (2022). Thirty years of resistance: Zig-zag through the plant immune system. Plant Cell 34, 1447–1478.

62. Ngou, B.P.M., Ahn, H.-K., Ding, P., and Jones, J.D.G. (2021). Mutual potentiation of plant immunity by cell-surface and intracellular receptors. Nature 592, 110–115.

63. Nomoto, M., Skelly, M.J., Itaya, T., Mori, T., Suzuki, T., Matsushita, T., Tokizawa, M., Kuwata, K., Mori, H., Yamamoto, Y.Y., et al. (2021). Suppression of MYC transcription activators by the immune cofactor NPR1 fine-tunes plant immune responses. Cell Rep 37, 110125.

64. Nomura, K., Melotto, M., and He, S.Y. (2005). Suppression of host defense in compatible plant-*Pseudomonas syringae* interactions. Curr Opin Plant Biol 8, 361–368.

65. Nomura, K., Debroy, S., Lee, Y.H., Pumplin, N., Jones, J., and He, S.Y. (2006). A bacterial virulence protein suppresses host innate immunity to cause plant disease. Science 313, 220–223.

66. Olate, E., Jiménez-Gómez, J.M., Holuigue, L., and Salinas, J. (2018). NPR1 mediates a novel regulatory pathway in cold acclimation by interacting with HSFA1 factors. Nat Plants 4, 811–823.

67. Orvar, B.L., Sangwan, V., Omann, F., and Dhindsa, R.S. (2000). Early steps in cold sensing by plant cells: the role of actin cytoskeleton and membrane fluidity. Plant J 23, 785–794.

68. Peng, Y., Yang, J., Li, X., and Zhang, Y. (2021). Salicylic acid: biosynthesis and signaling. Annu Rev Plant Biol 72, 761–791.

69. Rekhter, D., Ludke, D., Ding, Y., Feussner, K., Zienkiewicz, K., Lipka, V., Wiermer, M., Zhang, Y., and Feussner, I. (2019). Isochorismate-derived biosynthesis of the plant stress hormone salicylic acid. Science 365, 498–502.

70. Saleh, A., Withers, J., Mohan, R., Marqués, J., Gu, Y., Yan, S., Zavaliev, R., Nomoto, M., Tada, Y., and Dong, X. (2015). Posttranslational modifications of the master transcriptional regulator NPR1 enable dynamic but tight control of plant immune responses. Cell Host Microbe 18, 169–182.

71. Sarkar, S., Das, A., Khandagale, P., Maiti, I.B., Chattopadhyay, S., and Dey, N. (2018). Interaction of Arabidopsis TGA3 and WRKY53 transcription factors on *Cestrum yellow leaf curling virus* (CmYLCV) promoter mediates salicylic acid-dependent gene expression in planta. Planta 247, 181–199.

72. Seo, P.J., Kim, M.J., Park, J.Y., Kim, S.Y., Jeon, J., Lee, Y.H., Kim, J., and Park, C.M. (2010). Cold activation of a plasma membrane-tethered NAC transcription factor induces a pathogen resistance response in Arabidopsis. Plant J 61, 661–671.

73. Sheen, J. (2001). Signal transduction in maize and Arabidopsis mesophyll protoplasts. Plant Physiol 127, 1466–1475.

74. Spoel, S.H., and Dong, X. (2012). How do plants achieve immunity? Defence without specialized immune cells. Nat Rev Immunol 12, 89–100.

75. Stockinger, E.J., Gilmour, S.J., and Thomashow, M.F. (1997). *Arabidopsis thaliana CBF1* encodes an AP2 domain-containing transcriptional activator that binds to the C-repeat/DRE, a *cis*-acting DNA regulatory element that stimulates transcription in response to low temperature and water deficit. Proc Natl Acad Sci USA 94, 1035–1040.

76. Tada, Y., Spoel, S.H., Pajerowska-Mukhtar, K., Mou, Z., Song, J., Wang, C., Zuo, J., and Dong, X. (2008). Plant immunity requires conformational changes of NPR1 via S-nitrosylation and thioredoxins. Science 321, 952–956.

77. Tang, K., Zhao, L., Ren, Y., Yang, S., Zhu, J.K., and Zhao, C. (2020). The transcription factor ICE1 functions in cold stress response by binding to the promoters of *CBF* and *COR* genes. J Integr Plant Biol 62, 258–263.

78. Torrens-Spence, M.P., Bobokalonova, A., Carballo, V., Glinkerman, C.M., Pluskal, T., Shen, A., and Weng, J.K. (2019). PBS3 and EPS1 complete salicylic acid biosynthesis from isochorismate in Arabidopsis. Mol Plant 12, 1577–1586.

79. Vlot, A.C., Dempsey, D.A., and Klessig, D.F. (2009). Salicylic Acid, a multifaceted hormone to combat disease. Annu Rev Phytopathol 47, 177–206.

80. Wang, C., Dai, S., Zhang, Z.L., Lao, W., Wang, R., Meng, X., and Zhou, X. (2021). Ethylene and salicylic acid synergistically accelerate leaf senescence in Arabidopsis. J Integr Plant Biol 63, 828–833.

81. Wang, D., Amornsiripanitch, N., and Dong, X. (2006). A genomic approach to identify regulatory nodes in the transcriptional network of systemic acquired resistance in plants. PLoS Pathog 2, e123.

82. Wang, J., Hu, M., Wang, J., Qi, J., Han, Z., Wang, G., Qi, Y., Wang, H.W., Zhou, J.M., and Chai, J. (2019). Reconstitution and structure of a plant NLR resistosome conferring immunity. Science 364, 44–54.

83. Wang, W., Withers, J., Li, H., Zwack, P.J., Rusnac, D.V., Shi, H., Liu, L., Yan, S., Hinds, T.R., Guttman, M., Dong, X., and Zheng, N. (2020). Structural basis of salicylic acid perception by Arabidopsis NPR proteins. Nature 586, 311–316.

84. Wang, Y., Bao, Z., Zhu, Y., and Hua, J. (2009). Analysis of temperature modulation of plant defense against biotrophic microbes. Mol Plant Microbe Interact 22, 498–506.

85. Wang, Y., Pruitt, R.N., Nürnberger, T., and Wang, Y. (2022). Evasion of plant immunity by microbial pathogens. Nat Rev Microbiol 20, 449–464.

86. Wigge, P.A. (2013). Ambient temperature signalling in plants. Curr Opin Plant Biol 16, 661–666.

87. Wu, C.H., and Derevnina, L. (2023). The battle within: How pathogen effectors suppress NLR-mediated immunity. Curr Opin Plant Biol 74, 102396.

88. Wu, Z., Han, S., Zhou, H., Tuang, Z.K., Wang, Y., Jin, Y., Shi, H., and Yang, W. (2019). Cold stress activates disease resistance in *Arabidopsis thaliana* through a salicylic acid dependent pathway. Plant Cell Environ 42, 2645–2663.

89. Xin, X.F., Nomura, K., Aung, K., Velasquez, A.C., Yao, J., Boutrot, F., Chang, J.H., Zipfel, C., and He, S.Y. (2016). Bacteria establish an aqueous living space in plants crucial for virulence. Nature 539, 524–529.

90. Yang, H., Shi, Y., Liu, J., Guo, L., Zhang, X., and Yang, S. (2010). A mutant CHS3 protein with TIR-NB-LRR-LIM domains modulates growth, cell death and freezing tolerance in a temperature-dependent manner in Arabidopsis. Plant J 63, 283–296.

91. Yang, S., and Hua, J. (2004). A haplotype-specific resistance gene regulated by BONZAI1 mediates temperature-dependent growth control in Arabidopsis. Plant Cell 16, 1060–1071.

92. Ye, K., Li, H., Ding, Y., Shi, Y., Song, C., Gong, Z., and Yang, S. (2019). BRASSINOSTEROID-INSENSITIVE2 negatively regulates the stability of transcription factor ICE1 in response to cold stress in Arabidopsis. Plant Cell 31, 2682–2696.

93. Yu, X., Xu, Y., and Yan, S. (2021). Salicylic acid and ethylene coordinately promote leaf senescence. J Integr Plant Biol 63, 823–827.

94. Yu, X., Cui, X., Wu, C., Shi, S., and Yan, S. (2022). Salicylic acid inhibits gibberellin signaling through receptor interactions. Mol Plant 15, 1759–1771.

95. Yuan, M., Jiang, Z., Bi, G., Nomura, K., Liu, M., Wang, Y., Cai, B., Zhou, J.-M., He, S.Y., and Xin, X.-F. (2021). Pattern-recognition receptors are required for NLR-mediated plant immunity. Nature 592, 105–109.

96. Zavaliev, R., Mohan, R., Chen, T., and Dong, X. (2020). Formation of NPR1 condensates promotes cell survival during the plant immune response. Cell 182, 1093–1108.e1018.

97. Zeier, J. (2021). Metabolic regulation of systemic acquired resistance. Curr Opin Plant Biol 62, 102050.

98. Zhang, Y., Fan, W., Kinkema, M., Li, X., and Dong, X. (1999). Interaction of NPR1 with basic leucine zipper protein transcription factors that bind sequences required for salicylic acid induction of the *PR-1* gene. Proc Natl Acad Sci USA 96, 6523.

99. Zhao, C., Wang, P., Si, T., Hsu, C.C., Wang, L., Zayed, O., Yu, Z., Zhu, Y., Dong, J., Tao, W.A., et al. (2017). MAP kinase cascades regulate the cold response by modulating ICE1 protein stability. Dev Cell 43, 618–629.e615.

100. Zhou, J.M., Trifa, Y., Silva, H., Pontier, D., Lam, E., Shah, J., and Klessig, D.F. (2000). NPR1 differentially interacts with members of the TGA/OBF family of transcription factors that bind an element of the *PR-1* gene required for induction by salicylic acid. Mol Plant Microbe Interact 13, 191–202.

