## Supplemental Information-Version 2 for "Inducer of CBF Expression 1 (ICE1) Promotes Cold-enhanced Immunity by Directly Activating Salicylic Acid Signaling"

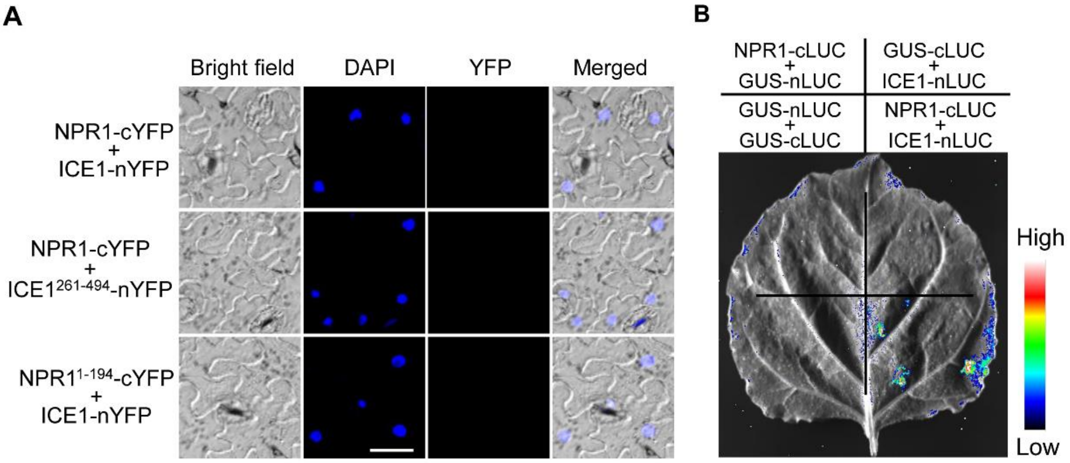


**Supplemental Fig. S1, supports Fig. 2.** No interacting signal of ICE1 and NPR1 was detected at normal temperature. (A) BiFC assay. Fluorescence was not observed in the nucleus of transformed *N. benthamiana* cells co-expressing ICE1-nYFP (or ICE1$^{261\text{-}494}$-nYFP) with NPR1-cYFP or ICE1-nYFP with NPR1$^{1\text{-}194}$-cYFP at normal temperature (22°C). Nuclei are indicated by DAPI staining. Scale bar = 20 μm. (B) LUC assay. ICE1-nLUC and NPR1-cLUC were co-expressed in *N. benthamiana* leaves, no luminescence intensity was detected at normal temperature (22°C). GUS-nLUC and GUS-cLUC were set as negative control.

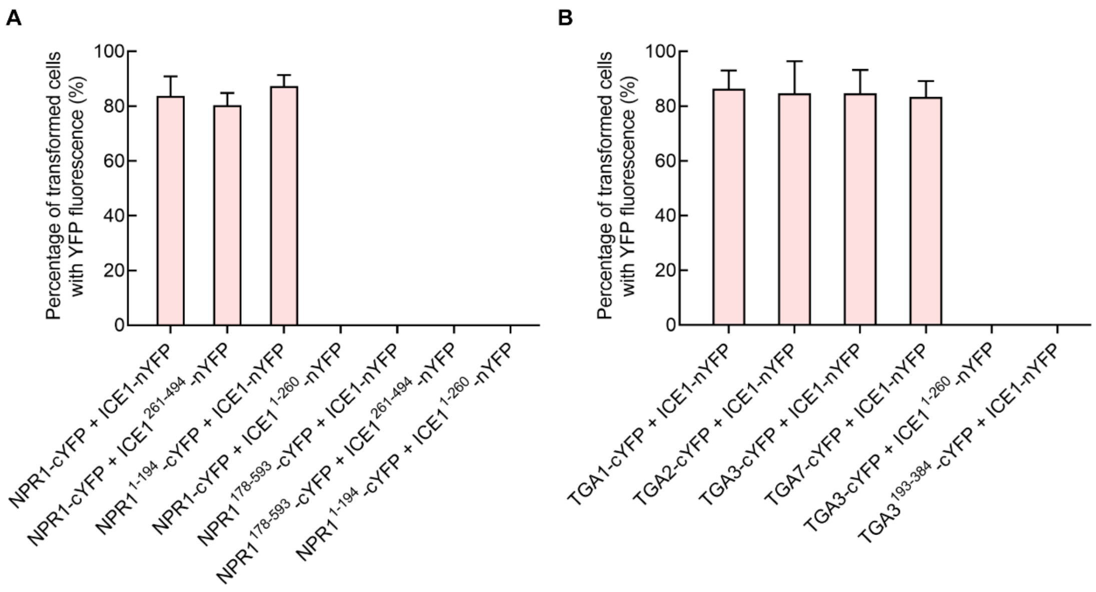


**Supplemental Fig. S2, supports Fig. 2B and 6B.** Percentage of transformed cells with YFP fluorescence in the BiFC assays presented in Fig. 2B and 6B. All experiments were performed three times, each evaluating more than 600 transformed cells. Values shown are mean ± s.d.

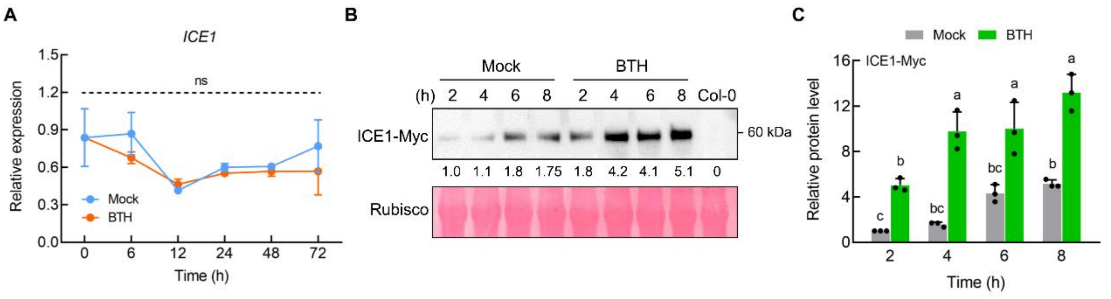


**Supplemental Fig. S3, supports Figure 3.** Transcriptional and protein levels of ICE1 responding to BTH treatment. (A) RT-qPCR analysis of *ICE1* expression in Col-0 leaves treated with or without 100 μM BTH in the indicating time before RNA extraction. Values are displayed as mean ± s.e.m. ($n$ = 3 biological replicates). (B) Immunoblot analyzing the accumulation of ICE1 protein induced by BTH. ICE1-overexpressing (*ICE1-Myc*) plants were treated with mock or 100 μM BTH for the indicated time before protein extraction, Col-0 used as negative control. The accumulation of ICE1 protein was detected with anti-Myc antibody. (C) Relative protein levels of ICE1-Myc in (B), which were quantified by Image J. Values are displayed as mean ± s.e.m. ($n$ = 3 biological replicates). Different letters indicate statistically significant differences (two-way ANOVA, $P < 0.05$).

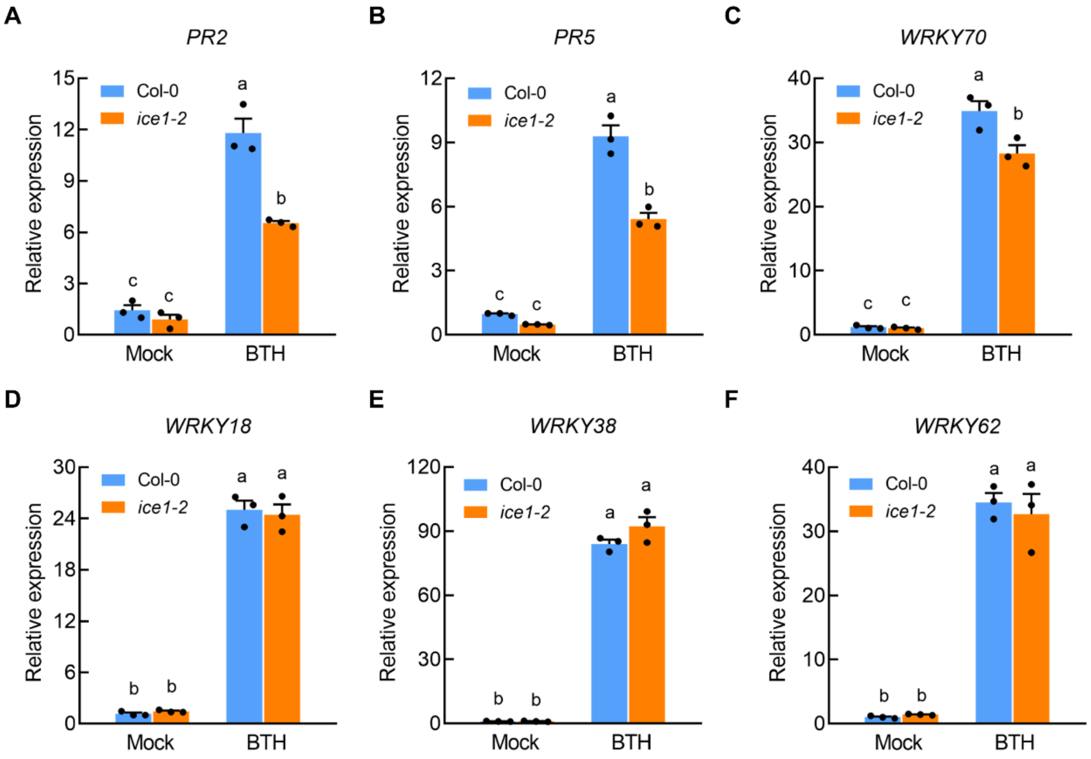


**Supplemental Fig. S4, supports Fig. 3.** RT-qPCR analysis of SA-related defense genes. (A-F) Testing the transcription levels of *PR2* (A), *PR5* (B), *WRKY70* (C), *WRKY18* (D), *WRKY38* (E) and *WRKY62* (F) in Col-0 and *ice1-2* treated with BTH, mock treatment used as control. Values are displayed as mean ± s.e.m. ($n$ = 3 biological replicates). Different letters indicate statistically significant differences (two-way ANOVA, $P < 0.05$). Experiments were repeated at least three times with similar trends.

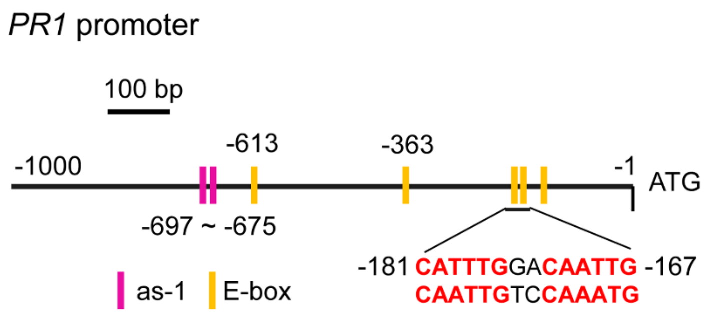


**Supplemental Fig. S5, supports Fig. 3G.** Diagram of promoter region of the *PR1* gene. Yellow boxes on the line indicate the potential ICE1 binding sites (MYC recognized sequences, called E-box). Magenta boxes on the line represent the *cis* elements (*as-1* elements, recognized by TGA proteins). The two E-box located in the area of -181 ~ -167 bp are bind by ICE1 in the EMSA assay.

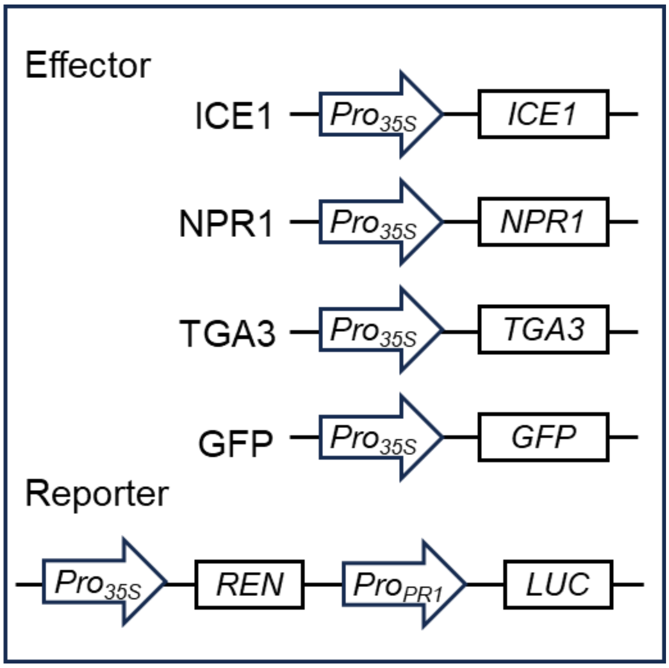


**Supplemental Fig. S6, supports Fig. 4 and Fig. 7.** Schematic of effectors and reporters used in transient transactivation assays. The effectors consisted of GFP, NPR1, TGA3 and ICE1 under the control of *Pro35S*, and the reporter consisted of the *PR1* promoter fused to the *LUC* gene.

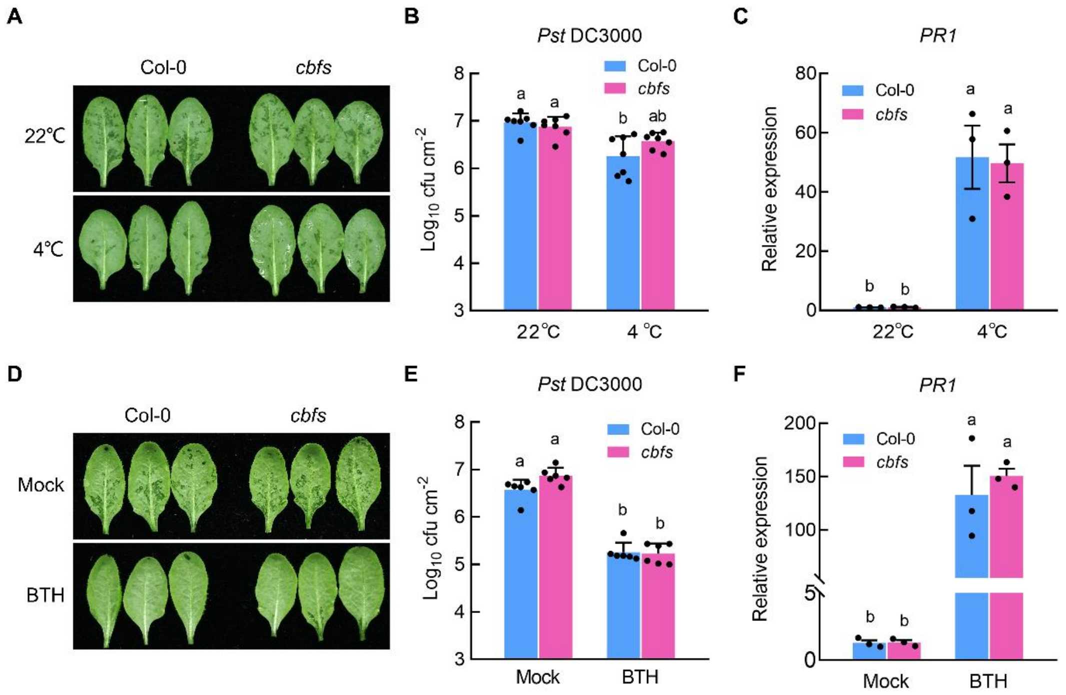


**Supplemental Fig. S7, supports Fig. 5.** *CBF* genes does not mediate low temperature-enhanced and SA-regulated immunity. (A) Col-0 and *cbfs* triple mutants leaves infiltrated with *Pst* DC3000 ($OD_{600}$ = 0.0001) after 10 hours of 4°C treatments (down), 22°C set as control (up). (B) Bacterial populations in leaves described in (A). Values are displayed as mean ± s.d. (*n* = 7 biological replicates). (C) RT-qPCR analysis of *PR1* expression in Col-0 and *cbfs* mutant leaves with or without 4°C treatment. Values are displayed as mean ± s.e.m. (*n* = 3 biological replicates). Experiments were repeated at least three times with similar trends. (D) Col-0 and *cbfs* triple mutants leaves infiltrated with *Pst* DC3000 ($OD_{600}$ = 0.0001) after 24 hours of pretreating with 100 μM BTH (down), mock set as control (up). (E) Bacterial populations in leaves described in (D). Values are displayed as mean ± s.d. (n = 6 biological replicates). (F) RT-qPCR analysis of *PR1* expression in Col-0 and *cbfs* mutant leaves with mock or BTH treatments. Values are displayed as mean ± s.e.m. (*n* = 3 biological replicates). Different letters indicate statistically significant differences (two-way ANOVA, $P < 0.05$). Experiments were repeated at least three times with similar trends.

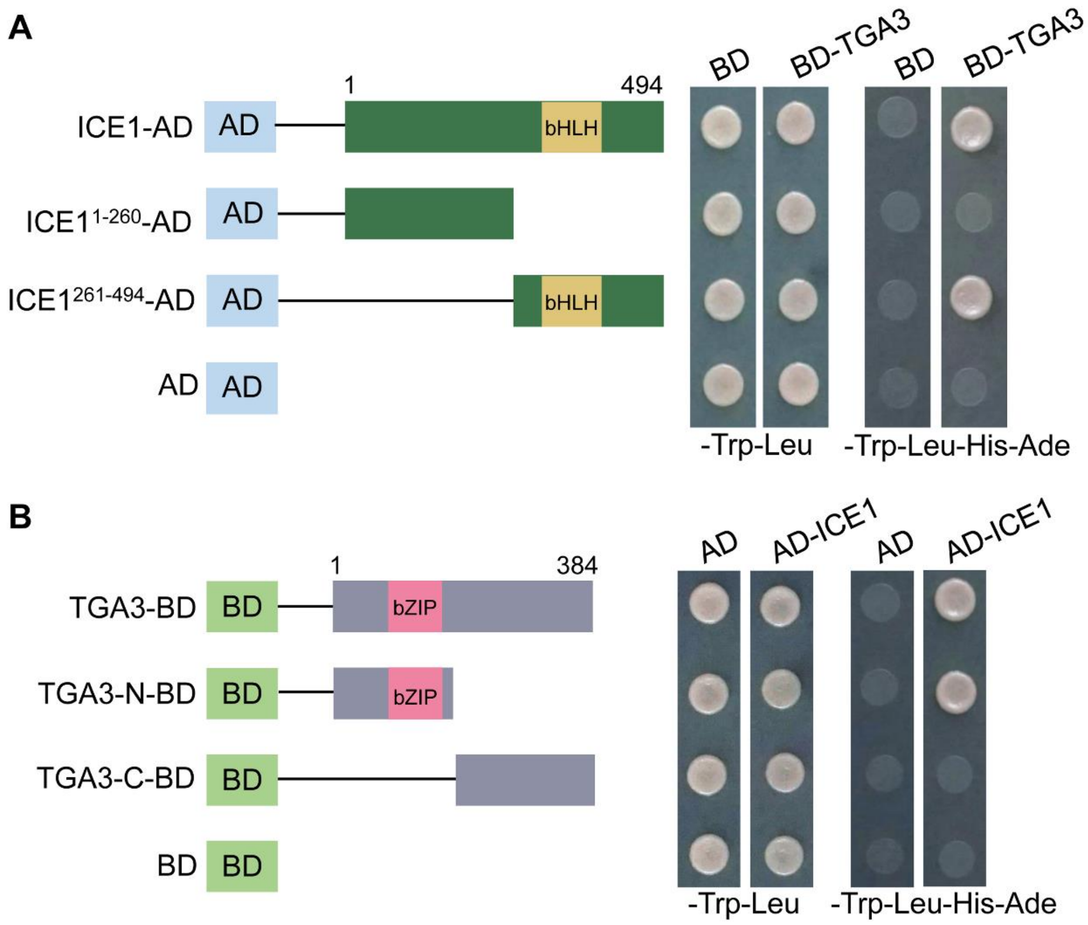


**Supplemental Fig. S8, supports Fig. 6.** The bHLH motif and bZIP motif was required for interaction of ICE1 and TGA3. (A) Yeast two hybrid assays showing the interactions between full-length and truncated ICE1 and TGA3. (B) Yeast two hybrid assays showing the interactions between full-length and truncated TGA3 and ICE1. Protein interactions were indicated by the growth of yeast cells after 2 days of incubation in dropout medium lacking Leu, Trp, His and Ade. The numbers indicate the positions of amino acids, pGBKT7 (BD) and pGADT7 (AD) were used as negative controls.

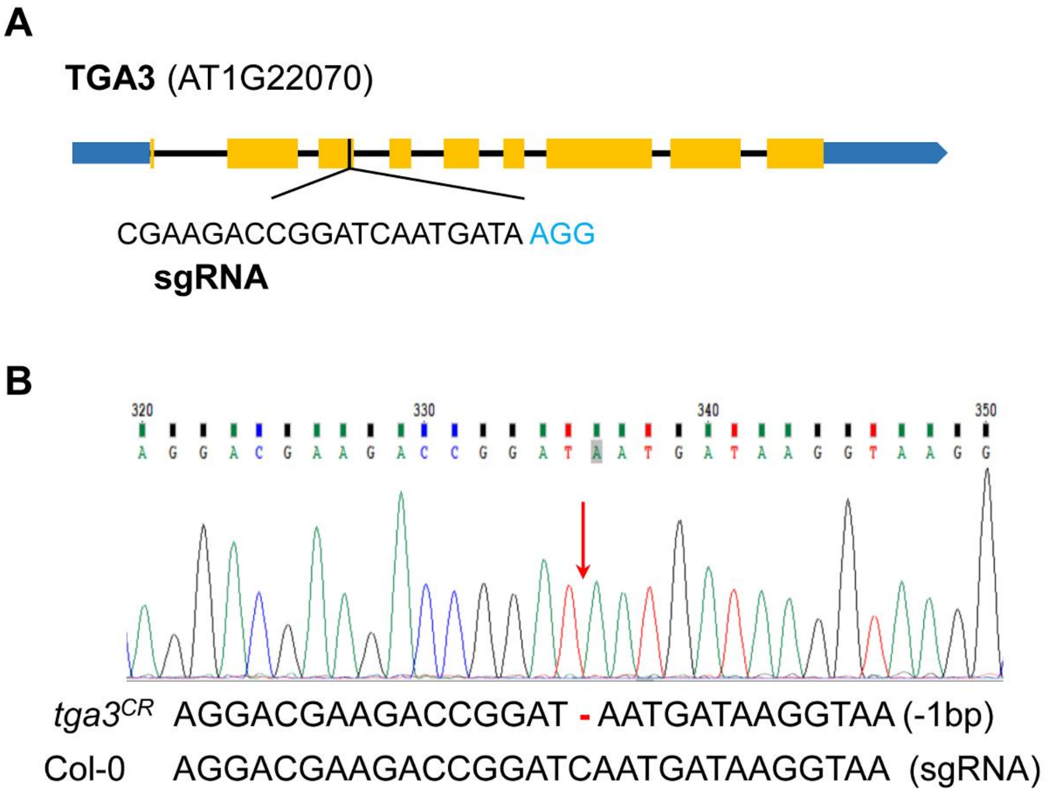


**Supplemental Fig. S9, supports Fig. 7.** Generation of *tga3*$^{CR}$ knockout by CRISPR/Cas9 technology. (A) The sgRNA targeting *TGA3* gene was designed to create *tga3*$^{CR}$. (B) One mutant, *tga3*$^{CR}$, was confirmed by Sanger sequencing and mutation type was indicated.

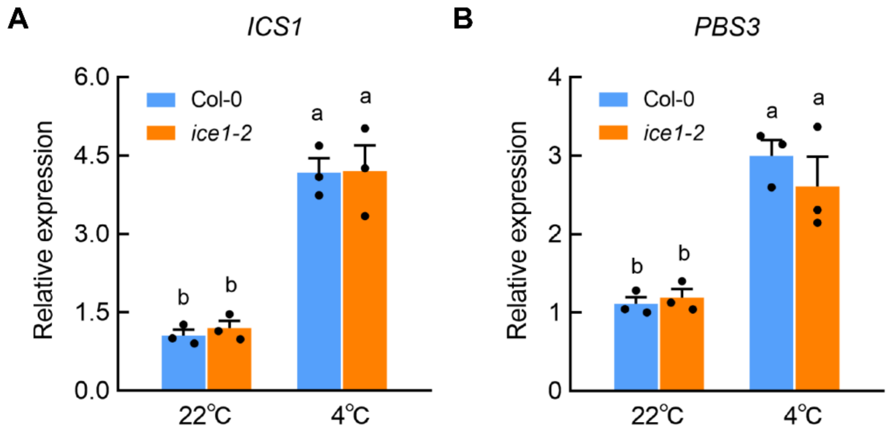


**Supplemental Fig. S10, supports Fig. 7.** RT-qPCR analysis of SA biosynthesis genes. (A-B) Transcription levels of *ICS1* (A), and *PBS3* (B) in Col-0 and *ice1-2* under normal and cold temperature conditions. Values are displayed as mean ± s.e.m. ($n$ = 3 biological replicates). Different letters indicate statistically significant differences (two-way ANOVA, $P < 0.05$). Experiments were repeated at least three times with similar trends.

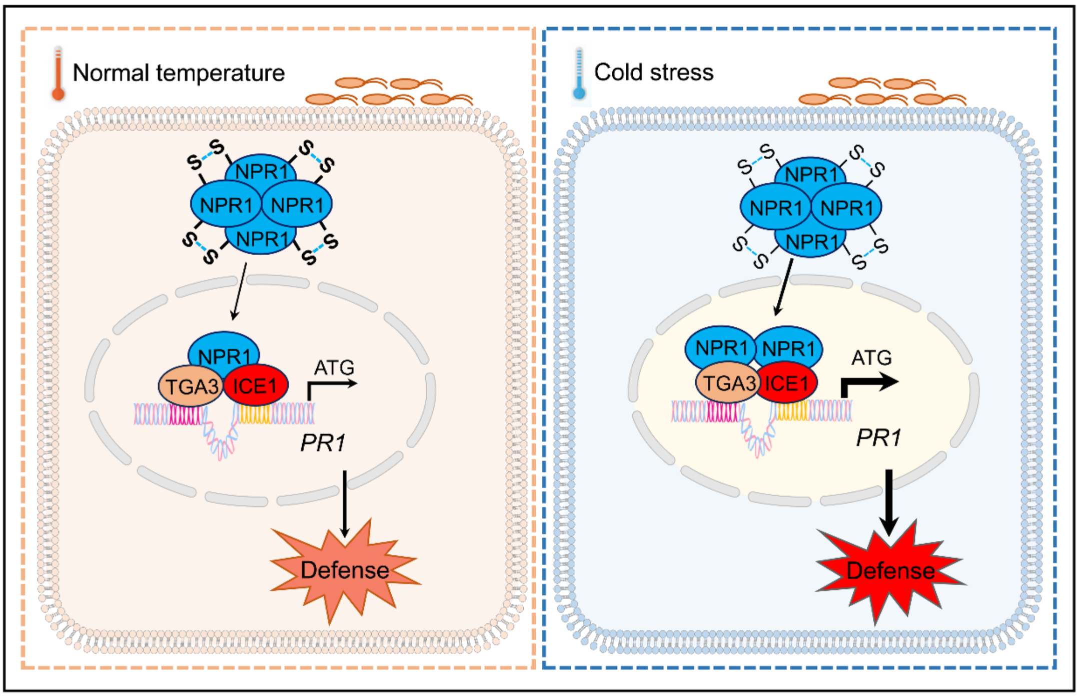


**Supplemental Fig. S11.** Working model of NPR1-TGA3/ICE1 regulating module against *Pst* DC3000. At normal temperature, NPR1 is depolymerized by high accumulation of SA caused by *Pst* DC3000 infection. Monomers of NPR1 are translocated into nucleus where it interacts with TGAs and ICE1 to activate their transcriptional activations on expression of pathogen-relative gene, *PR1*, to promote plant immune responses against pathogen attacks. Under low temperature, the cold-induced translocation of NPR1 enhances the activation of ICE1 and TGA3 on *PR1* expression, which causes robust plant defense response against pathogen attacks. In the promoter of *PR1*, the area from -613 to -137 bp contains five E-boxes and the two E-boxes located in the yellow area of -181 ~ -167 bp are bind by ICE1, while the magenta area from -697 to -675 has two *as-1* elements that bind by TGA3. The information of *PR1* promoter was obtained from the Arabidopsis Information Web site (https://www.arabidopsis.org/).

**Supplemental Fig. S12, supports Fig. 2D, Fig. 2E, Fig. 5E, Fig. 6D and Fig. S4B.** Uncropped images for protein gels.

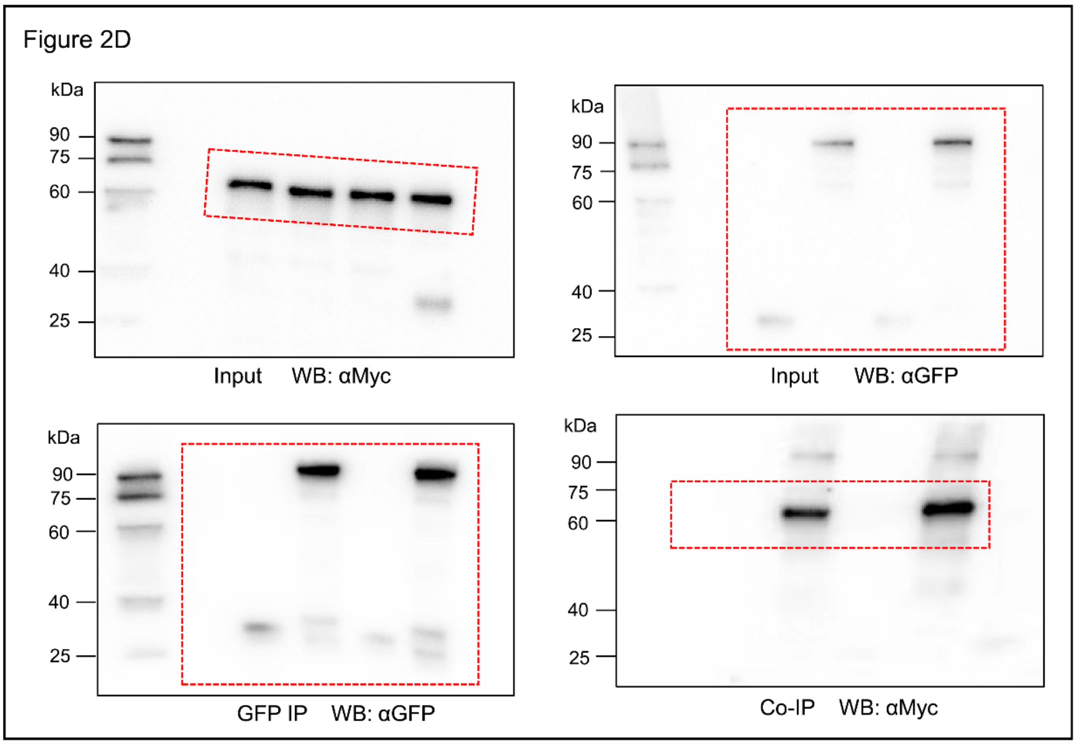


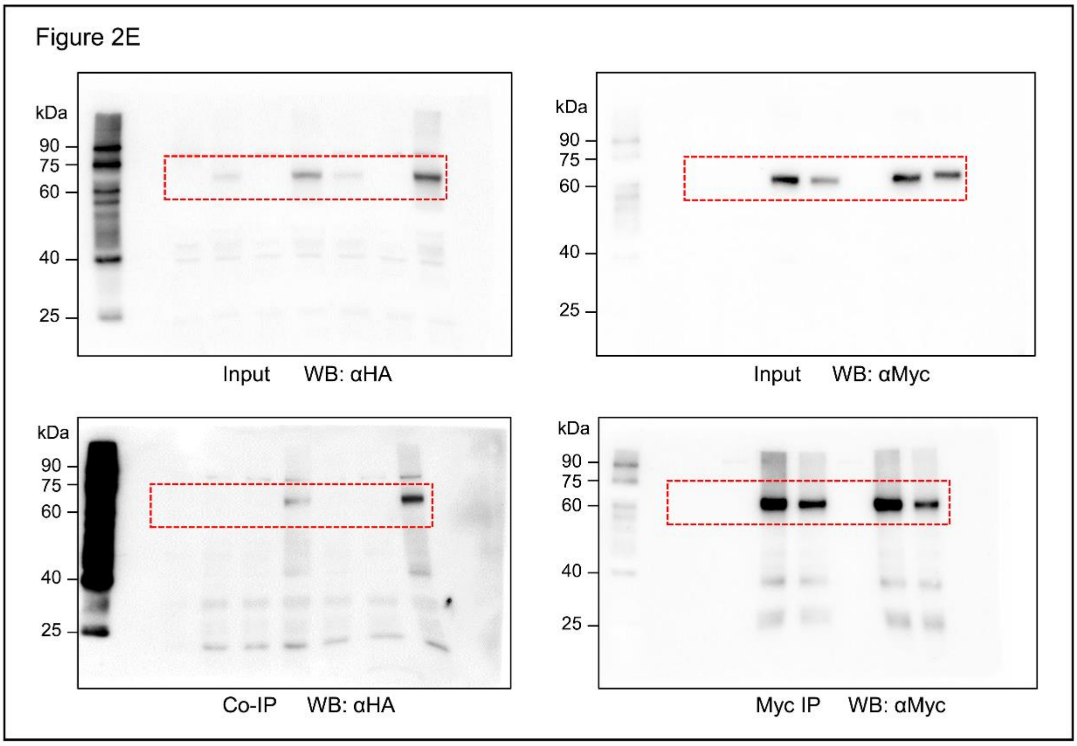

Figure 5E

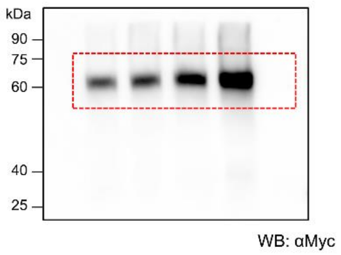


Figure S4B

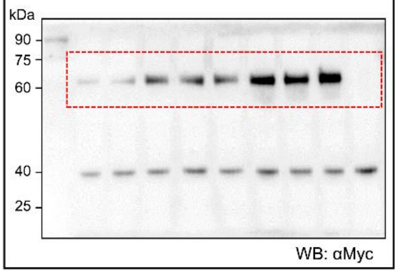


Figure 6D

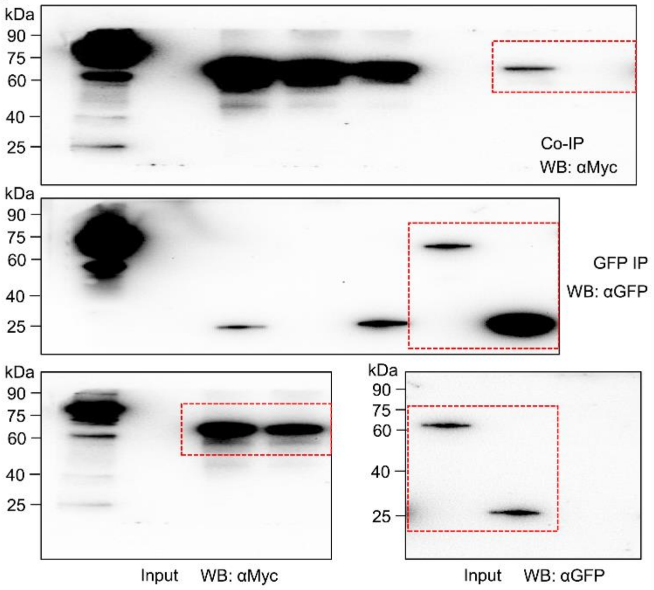
